# Discovery of a pathway-selective platelet P2Y_1_R inverse agonist that suppresses inflammation while preserving hemostasis

**DOI:** 10.64898/2026.06.19.732319

**Authors:** Simon C. Pitchford, Kazi Nahar, Dingxin Pan, Claudia M. Sisk, Taha Al-Adhami, Kubra Ekinci, Richard T. Amison, Nupur Gargate, Anjana Saji, Edward Wills, Clive P. Page, Graham Ladds, Khondaker Miraz Rahman

## Abstract

The platelet P2Y_1_ receptor (P2Y_1_R) is necessary for inflammation, signalling via Rho-GTPase pathways to elicit functions that are distinct from aggregation (PLC-dependent canonical signalling pathway). Whether these distinct platelet inflammatory functions can be selectively suppressed to preserve hemostasis through the rational design of P2Y_1_R antagonists has not been explored. *In silico* molecular docking analysis examined biased nucleotide interactions within the P2Y_1_R binding pocket. The identified possible key amino acid residues guided rational design to synthesize compounds for pathway selective inhibition, evolving from nucleotide to non-nucleotide structures. The nucleotide analogue KMR-82-13 was predicted to engage distinct regions of the binding pocket and selectively inhibited platelet chemotaxis while preserving aggregation. These findings informed the design of a non-nucleotide compound KSN-159-27, aiming to retain key KMR-82-13-like interactions while improving drug-like properties. Docking and molecular dynamics simulation supported a stable but dynamic binding mode for KSN-159-27 within the P2Y_1_R pocket, consistent with pathway-selective inhibition. KSN-159-27 displayed characteristics of a pathway selective inverse agonist at P2Y_1_R towards G_α12/13_-mediated pathways, but not those associated by G_αq_ activation in P2Y_1_R-transfected HEK293T cells. KSN-159-27 showed functionally selective inhibition for platelet P2Y_1_R-mediated functions. *In vivo*, KSN-159-27 suppressed inflammatory cell recruitment, whilst preserving bleeding time and ADP-induced thromboembolic responses, in contrast to the ‘neutral’ P2Y_1_R antagonist MRS2500. This first demonstration for the rational design of a pathway selective inverse agonist at platelet P2Y_1_Rs has significant implications for novel therapeutic strategies developed to safely target platelet activation during inflammation, in contrast to current anti-platelet drugs used in the prevention of thrombosis.

**Key Points:**

- Biased inverse platelet P2Y_1_R agonists selectively supress inflammation whilst preserving hemostasis and the ability of platelets to aggregate.
- Biased inverse agonism selectively inhibited P2Y_1_R G_α12/13_ (Rho-GTPAse functions) but not G_αq_ activities (PLC functions).

## Introduction

Platelet activation is essential for host defence and inflammation, providing a valid cellular target to develop novel anti-inflammatory drugs.^1,2^ However, platelets are best known for their requisite role during hemostasis and thrombosis which require distinct functions compared to inflammation.^3^ The activity of platelets during both hemostasis and inflammation is highly regulated by the surrounding nucleotide and nucleoside microenvironment, known as the ‘purinome’.^4–6^ Whilst it is not understood how distinct inflammatory and hemostatic platelet functions are physiologically regulated, they can nevertheless occur via P2Y_1_R activation.^7–12^ Platelet P2Y_1_R can be activated by various endogenous nucleotides in addition to the cognate ligand ADP to stimulate Rho-GTPase dependent platelet motility and low-level P-selectin expression.^13^ However, these nucleotides do not activate the P2Y_1_R phospholipase C-β1 (PLC) pathway to cause platelet aggregation.^13^ This agonist activity profile at P2Y_1_R can be explained via the concept of biased agonism.^14,15^ Molecularly, P2Y_1_R displays considerable G protein promiscuity, activating members of all four G protein families,^16^ in addition to signalling from β-arrestin recruitment.^17^ This diversity in protein-receptor coupling provides a plausible mechanism for biased agonism at P2Y_1_R, as opposed to an alternative system or environmental bias. A possible explanation for the contradiction in P2Y_1_R control of divergent platelet hemostatic and inflammatory functions is through changes in the purinome to differentially affect G_αq/11_ (PLC) and G_α12/13_ (RhoA/Rac1) activation.

Our objective was to better understand ligand docking within the P2Y_1_R orthosteric binding pocket,^13,18^ using an *in silico* approach to design and synthesize compounds for an assessment of their ability to selectively inhibit P2Y_1_R-dependent inflammatory functions of platelets, whilst preserving platelet hemostatic (aggregatory) functions.

## Methods

Detailed protocols, experimental design and statistical analysis are provided in the Supplementary Data (see *Blood* Website).

### Molecular docking, molecular dynamics simulation, and synthesis of P2Y**_1_**R ligands

Molecular docking was performed using the crystal structure of the human platelet P2Y_1_R (PDB ID: 4XNW). Chain A was prepared in Schrödinger Maestro 2025-4 using the Protein Preparation Wizard.^19^ Molecular dynamics simulation of the P2Y_1_R–KSN-159-27 complex was performed in Schrödinger Maestro/Desmond 2025-2 using the OPLS4 force field.^20^ Prime MM-GBSA binding free-energy analysis was then performed. The synthetic route for the ligands commenced from commercially available starting materials and progressed through a series of well-defined transformations.

### Human platelet studies

Blood was collected in accordance with local ethical approval from King’s College London (KCL, Research Ethics Committee reference: 10/H0807/99) and the Human Tissue Act 2004.^13^ Blood was processed to harvest platelets to measure aggregation, chemotaxis; and commercial assays to measure RhoA and Rac1 activity and IP1 production.

### Receptor pharmacology studies

HEK293T cells for signalling profiling were transfected with pcDNA3.1(+)/P2RY1 or pcDNA3.1(+)/P2RY12 from cDNA Resource Center (cDNA.org) and subjected to G-CASE G-protein dissociation assays,^21^ luminescence based genetic reporter assays,^22^ and intracellular Ca^2+^ mobilization assays.

### *In vivo* experimentation

Animal experiments were performed in accordance with the Animals (Scientific Procedures) Act 1986 with 2012 amendment, and local ethical approval. BALB/c mice were sourced from Charles River Laboratories Ltd. Littermate male and female test/platelet P2Y ^-/-^ mice and control ‘wild type’ mice were bred in house.^12^ Mice were used for LPS-induced pulmonary inflammation, and thrombo-embolism studies.

## Results

### *In silico* molecular docking analyses of P2Y_1_R ligands to predict interactions required for functionally-selective pathway specific inhibition

Utilizing molecular modelling approaches, we identified key residues (Val194^ECL2^, Lys196^ECL2^ and Thr205^ECL2^) within P2Y_1_R that were predicted to participate in the binding of functionally biased endogenous nucleotides (e.g. NAD^+^, Up4A and ADP-ribose) to form a spatially defined interaction region deeper within the orthosteric pocket that is absent in interactions with ADP.^13,18,23^ Molecular modelling of the ‘neutral’ P2Y_1_R antagonist MRS2500 (**Figure 1A-C**) within this P2Y_1_R binding pocket identified the iodine substitution on the purine ring as an optimal position for chemical modification to introduce interactions with residues associated with functionally biased nucleotides to generate a functionally selective antagonist scaffold. Guided by this structural basis, we designed a novel chemical structure, **KMR-82-13** (**Figure 1D**), by replacing the iodine substituent of MRS2500 with a heteroaliphatic functionality capable of engaging these residues while maintaining the overall steric and physicochemical characteristics required for binding within the orthosteric pocket. Multiple *in silico* modifications were evaluated, and docking analysis indicated that substitution of the iodine group with a methylpiperazine provided an optimal interaction profile. In 3D and 2D interaction analyses, this modification enabled favourable interactions with Val194^ECL2^, Lys196^ECL2^ and Thr205^ECL2^ residues (**Figure 1E and 1F**). This methylpiperazine extension also introduced a previously absent interaction with Gln307^7.36^ and Cys202^45.50^ which MRS2500 does not interact with (**Figure 1B and 1C**) but were key interaction residues for ADP and the endogenous biased nucleotides. ^13^ This interaction further stabilized the ligand deeper within the orthosteric cavity. Furthermore, like the endogenous biased nucleotides, but unlike ADP and MRS2500, **KMR-82-13** was predicted not to interact with Lys46^NTerm^ or Asn283^6.58^ (**Figure 1E and 1F**). The rational design of **KMR-82-13** therefore represented an antagonist scaffold predicted to selectively disrupt P2Y_1_R-dependent platelet inflammatory functions while preserving hemostatic responses. **KMR-82-13** was subsequently synthesized to validate the *in silico* predictions through experimental studies.

**Figure 1.**
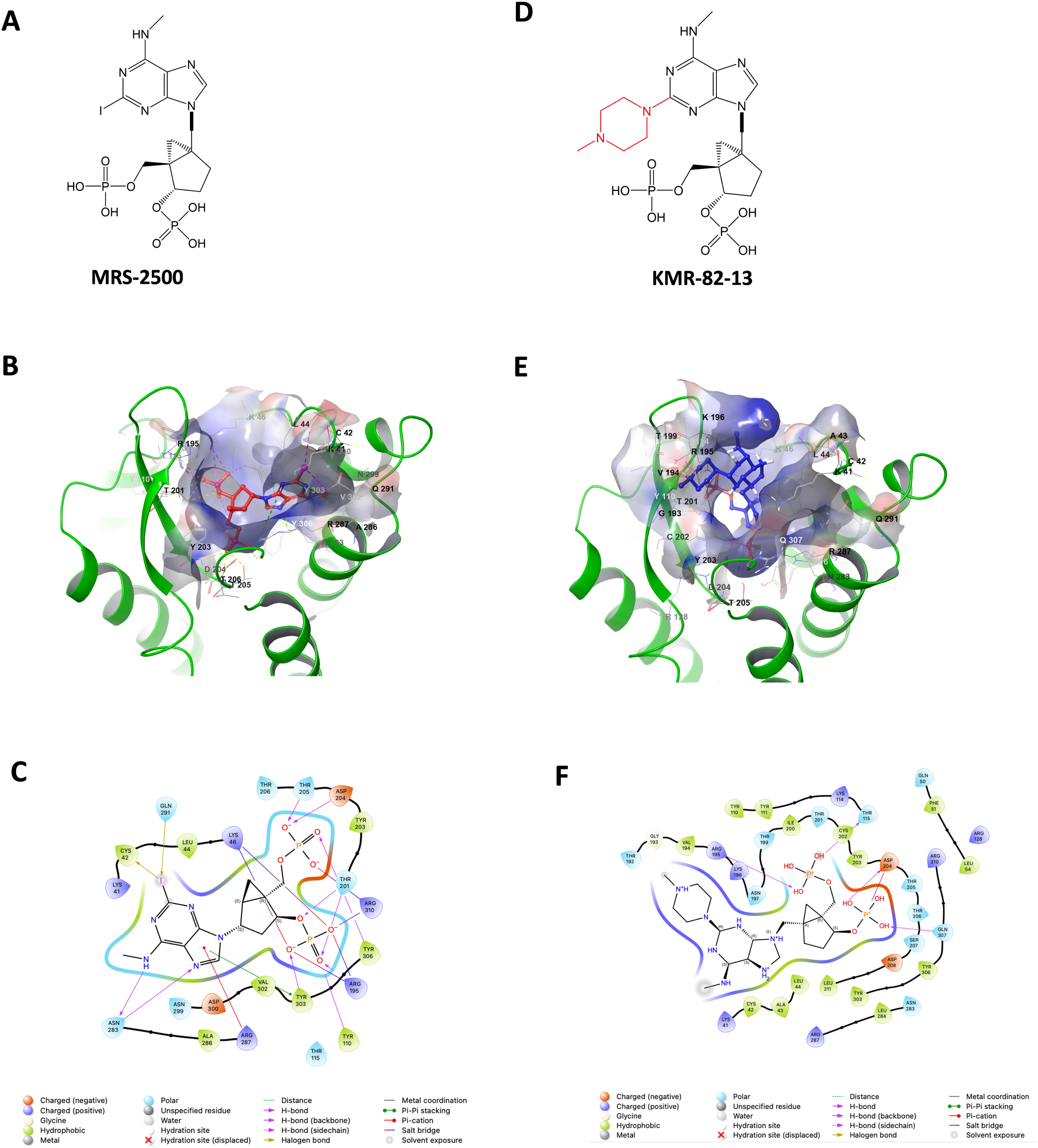
Molecular modelling of the binding pocket of the P2Y_1_ receptor to select structure KMR82-13. Structure of MRS2500 (**A**), and 3D and 2D image of binding pocket of MRS2500 (**B**, **C**). Structure of KMR-82-13 (**D**). The methylpiperazine substitution is shown in red. 3D and 2D image of binding pocket of KMR-82-13 (**E**, **F**) showing distinct interactions with residues within the P2Y_1_ receptor.

### Platelet P2Y_1_R activation can be selectively inhibited to supress inflammation whilst preserving aggregation and hemostasis

Defining characteristics of platelet P2Y_1_R-dependent functions during inflammation and infection include platelet migration, and platelet-dependent leukocyte recruitment.^7,24-28^ These functions can be mimicked using *in vitro* chemotaxis assays that are distinct from platelet aggregation and fibrinogen-binding.^13,18^ The incubation of platelets with **KMR-82-13** (from 10nM) inhibited chemotaxis towards fMLP after platelet activation with the selective P2Y_1_R agonist MRS2365 (**Figure 2A**) and also inhibited platelet-induced neutrophil chemotaxis (**Figure 2B**). The degree of inhibition was equivalent to when platelets were incubated with MRS2500 (**Figure 2A and 2B**). In contrast, platelet aggregation towards MRS2365 was not inhibited by **KMR-82-13** at any concentration tested up to 10µM, in contrast to MRS2500 (**Figure 2C and 2D**).

**Figure 2.**
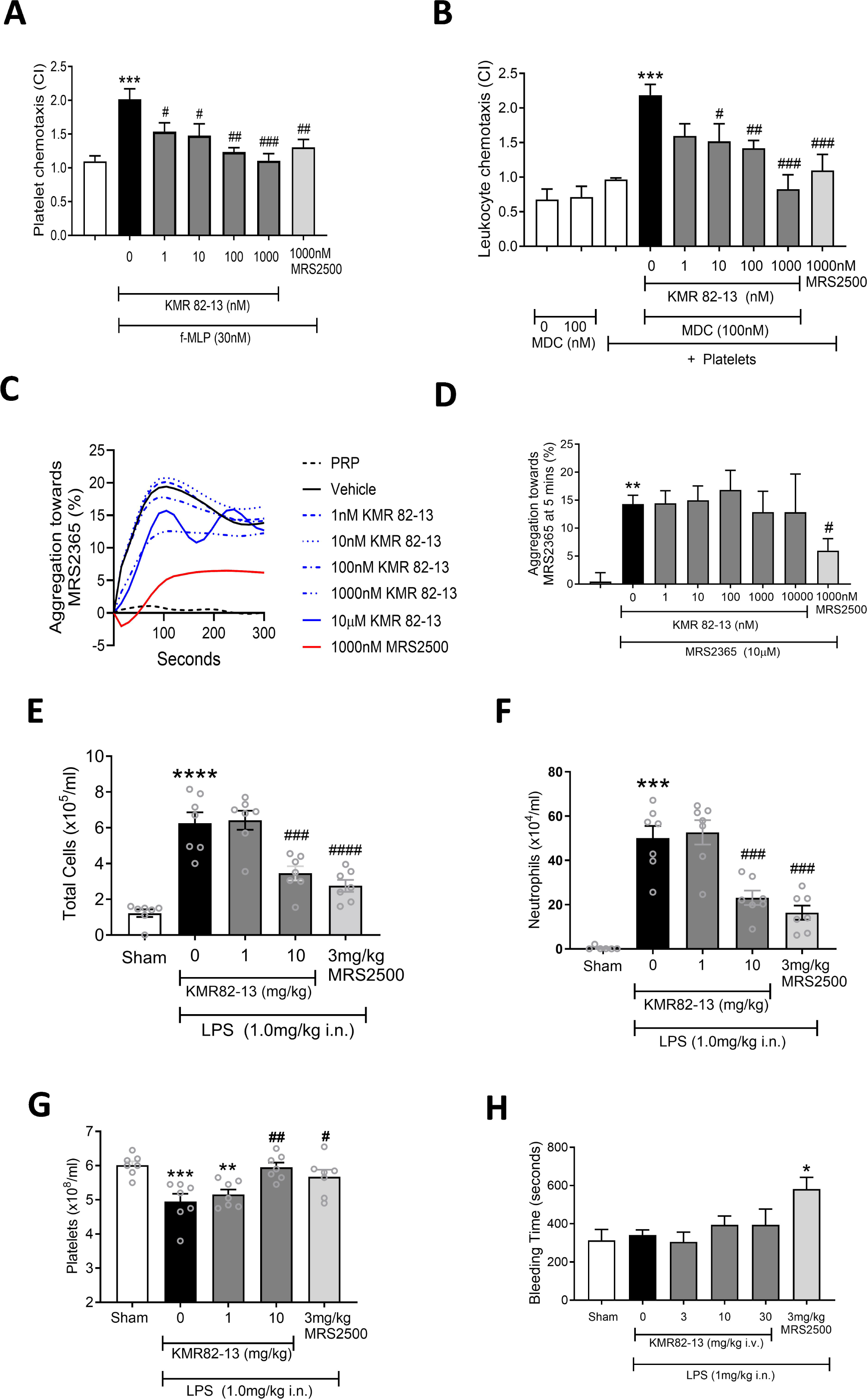
Effect of KMR-82-13 on P2Y_1_ dependent platelet inflammatory functions and P2Y_1_ induced aggregation *in vitro and in vivo*. Platelets from healthy human donors were incubated with KMR-82-13 or MRS2500 for 10 minutes before addition of 100nM MRS2365 (**A-D**) and subjected to chemotaxis towards 30nM fMLP (**A**), and platelet-dependent neutrophil chemotaxis towards 100nM MDC chemokine (**B**). In other experiments, platelet aggregation towards MRS2365 (10µM) was assessed (**C**), and analysed after 5 minutes (**D**). In other experiments, female Balb/c mice were administered KMR-82-13 or MRS2500 (intravenously), 5-10 minutes before intranasal administration of LPS (1mg/kg). 4 hours later, bronchoalveolar lavages were conducted to allow enumeration of total leukocyte counts (**E**), neutrophil counts (**F**). Tail snips were undertaken to allow measurement of circulating platelet numbers (**G**). A separate cohort of mice were utilized to measure tail bleeding time 60 minutes post LPS administration (**H**). n= 6 per group. Data means ±SEM. * P<0.05, *** P<0.001 vs white bar, # P<0.05, ## P<0.01, ### P<0.001 vs black bar.

Subjecting mice to LPS-induced lung inflammation, the administration of **KMR-82-13** (10mg/kg, i.v.) significantly inhibited the inflammatory response (**Figure 2E and 2F**), with equivalent efficacy to mice administered MRS2500 (3mg/kg, 65-70% inhibition). Circulating platelet numbers were maintained after **KMR-82-13** administration (**Figure 2G**). Tail bleeding times conducted as a measure of platelet-dependent hemostasis revealed that **KMR-82-13** (30mg/kg, i.v.) had no effect, in contrast to significant prolongation of bleeding with MRS2500 (**Figure 2H**). Thus, the design of a nucleotide structure based on predicted amino acid interactions of biased agonists was able to inhibit platelet P2Y_1_R-dependent inflammatory functions, whilst preserving hemostasis.

### Creation of non-nucleotide small molecule scaffolds to allow pharmacological characterization of pathway-specific, functionally selective P2Y_1_R inhibition

**KMR-82-13** is structurally derived from MRS2500, which loses potency rapidly following intravenous administration with no detectable effect at 60 mins.^29^ Whilst increasing the dose can improve the duration of action, this is not ideal for comparisons of potential dose related effects of distinct functions that will be constrained by dose limiting administration *in vivo*. Knowledge of the predicted P2Y_1_R orthosteric binding pocket interactions of **KMR-82-13** provided an opportunity to design non-nucleotidic small molecules capable of reproducing the same key interaction network within the P2Y_1_R binding pocket. Such scaffolds offer greater flexibility and diversification for medicinal chemistry optimization, including modulation of pharmacological characterization of receptor affinity, selectivity, and pharmacokinetic properties. During this process we examined previously reported ligands with P2Y_1_ inhibitory activity and identified the non-nucleotide compound **5**, which had been reported to possess oral bioavailability but without mechanistic information describing its binding mode.^30^

We designed **KSN-159-27** (**Figure 3A**) using the chemical scaffold of compound **5** as the template, with the aim to retain the key receptor interactions and binding profile observed for the biased interaction profile of **KMR-82-13**. **KSN-159-27** occupies the same deep orthosteric binding pocket as **KMR-82-13** (**Figure 3B-3D**), rather than the more superficial occupancy in this region engaged by MRS2500 (**Figure S1**). Importantly, the predicted orientation of **KSN-159-27** within the receptor aligned with the region of the pocket defined by interactions with Val194^ECL2^, Lys196^ECL2^ and Thr205^ECL2^ and neighbouring residues.

**Figure 3.**
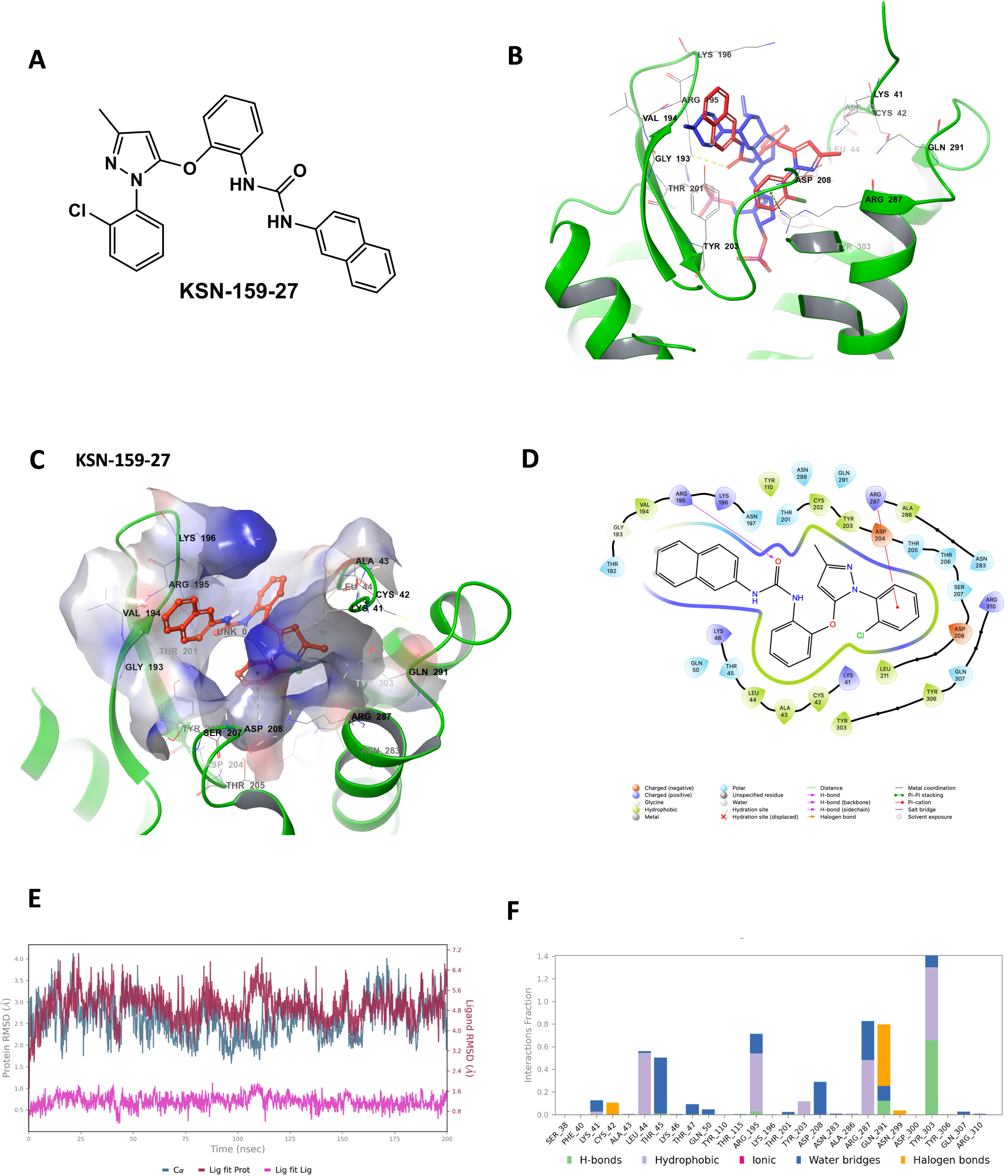
Design principles, docking analysis and molecular dynamics validation of KSN-159-27. (**A**) Chemical structure of KSN-159-27. (**B**) Overlay of the predicted docking poses of KMR-82-13 and KSN-159-27 within the crystal structure of P2Y_1_R, showing that both ligands occupy a similar binding site of the orthosteric binding pocket. (**C**,**D**) Three-dimensional and two-dimensional interaction maps of KSN-159-27 within the P2Y_1_R orthosteric binding site. (**E**) Protein–ligand RMSD analysis of the P2Y_1_R–KSN-159-27 complex during the 200 ns membrane molecular dynamics simulation. (**F**) Protein–ligand contact profile of the P2Y_1_R–KSN-159-27 complex during the 200 ns simulation

To assess whether this binding mode was dynamically retained, we performed a 200 ns membrane molecular dynamics simulation of the P2Y_1_R–KSN-159-27 complex. After equilibration, the receptor remained structurally stable, with a Cα RMSD of 2.61± 0.44 Å and no evidence of progressive structural drift. **KSN-159-27** showed local positional adaptation within the binding pocket, but its internal conformation remained stable, as reflected by a low ligand-fitted RMSD of 1.19± 0.24 Å and a stable radius of gyration (**Figure 3E**).

Protein RMSF analysis (**Figure S2A**) further supported stability of the ligand-binding region. The largest fluctuations were observed mainly in flexible loop or terminal regions outside the principal ligand-contacting pocket, including residues around 215–259 and the C-terminal region. In contrast, several residues contributing to ligand contacts showed lower Cα RMSF values, including Leu44^NTerm^, Thr45^NTerm^, Tyr203^45.51^, Asp208^ECL2^, Arg287^6.62^, Gln291^ECL3^, Tyr303^7.32^, Tyr306^7.35^, Gln307^7.36^ and Arg310^7.39^. This suggests that the binding pocket remained comparatively ordered while distal receptor regions retained expected flexibility.

Protein–ligand contact analysis showed **KSN-159-27** retained a recurrent interaction network within the P2Y_1_R binding cavity (**Figure 3F**). The most persistent contacts involved Tyr303^7.32^, Gln291^ECL3^, Arg287^6.62^ and Arg195^ECL2^, with additional contributions from Leu44^NTerm^, Thr45^NTerm^ and Asp208^ECL2^. These interactions indicate **KSN-159-27** did not remain fixed in a single static docking pose but instead underwent local adaptation while remaining accommodated within the same predicted binding region. Full RMSF and secondary-structure analyses are provided (**Figure S2**).

To further assess the energetic stability of the predicted binding mode, Prime MM-GBSA analysis was performed using 91 representative frames extracted every 2 ns from the equilibrated portion of the 200 ns trajectory. The P2Y_1_R–KSN-159-27 complex showed a favourable mean binding free energy of −60.73± 7.88 kcal/mol, supporting stable accommodation of **KSN-159-27** within the receptor binding pocket during the simulation. The binding energy was mainly supported by van der Waals (−46.08± 3.54 kcal/mol), lipophilic (−19.31± 1.36 kcal/mol) and Coulombic (−16.09± 5.01 kcal/mol) contributions. The polar solvation term opposed binding (23.94± 5.17 kcal/mol), consistent with the expected energetic cost of ligand desolvation upon complex formation. These energetic components align with the MD interaction profile, where **KSN-159-27** maintained recurrent hydrophobic, aromatic/cation-associated, polar and water-mediated contacts within the P2Y_1_R binding pocket.

Together, the docking, MD and MM-GBSA results indicate that **KSN-159-27** formed a dynamically stable and energetically favourable complex with P2Y_1_R. This structural model guided the design of a broader non-nucleotide ligand series with improved affinity and physicochemical properties, while preserving the key interaction pattern associated with **KMR-82-13**. The design strategy therefore focused on placing the compound **5**-derived core scaffold within the deeper orthosteric cavity and orienting substituents to reproduce the critical interactions identified for **KMR-82-13**. Using scaffold-hopping approaches combined with iterative *in silico* docking (**Figure S3**), we designed and synthesized focused KSN, TA and SL series of compounds to define the structure–activity relationship contributing to pathway-specific P2Y_1_R inhibition (Supplemental data: Chemistry section, Scheme 1).

### Comparison of P2Y_1_R cellular responses and platelet functions across a chemical series

We investigated whether there was a relationship within the compound library using cell-based pharmacological signalling assays and their ability to antagonise P2Y_1_R G_αq_-PLC (reporter NFAT-RE), G_βγ_-Raf (reporter SRE), and G_α12/13_-RhoA (reporter SRF-RE) -dependent activities (**Figure 4A**).^22^ MRS2500 showed similar potency at inhibiting each P2Y_1_R activity (**Figure 4B**), whilst compounds **KSN-159-27**, **KSN-159-28**, **TA-167-129**, **TA-167-153** and **TA-167-163** showed selective antagonism of SRF-RE and SRE-RE activities compared to NFAT-RE (**Figure 4B**). Compounds **TA-167-137**, **TA-167-139**, **TA-167-157**, **TA-167-161**, and **SL-183-5** revealed no antagonism for each of the P2Y_1_R activities (**Figure 4B**). Furthermore, all compounds failed to inhibit signalling activities in HEK293T cells transfected with P2Y_12_R (**Figure 4C**), providing evidence of compound specificity to purinergic receptors activated by ADP on platelets. In these assays, AR-C66096 (P2Y_12_R antagonist^4^) was the only compound to display significant antagonism of P2Y_12_R-dependent signalling activity.

**Figure 4.**
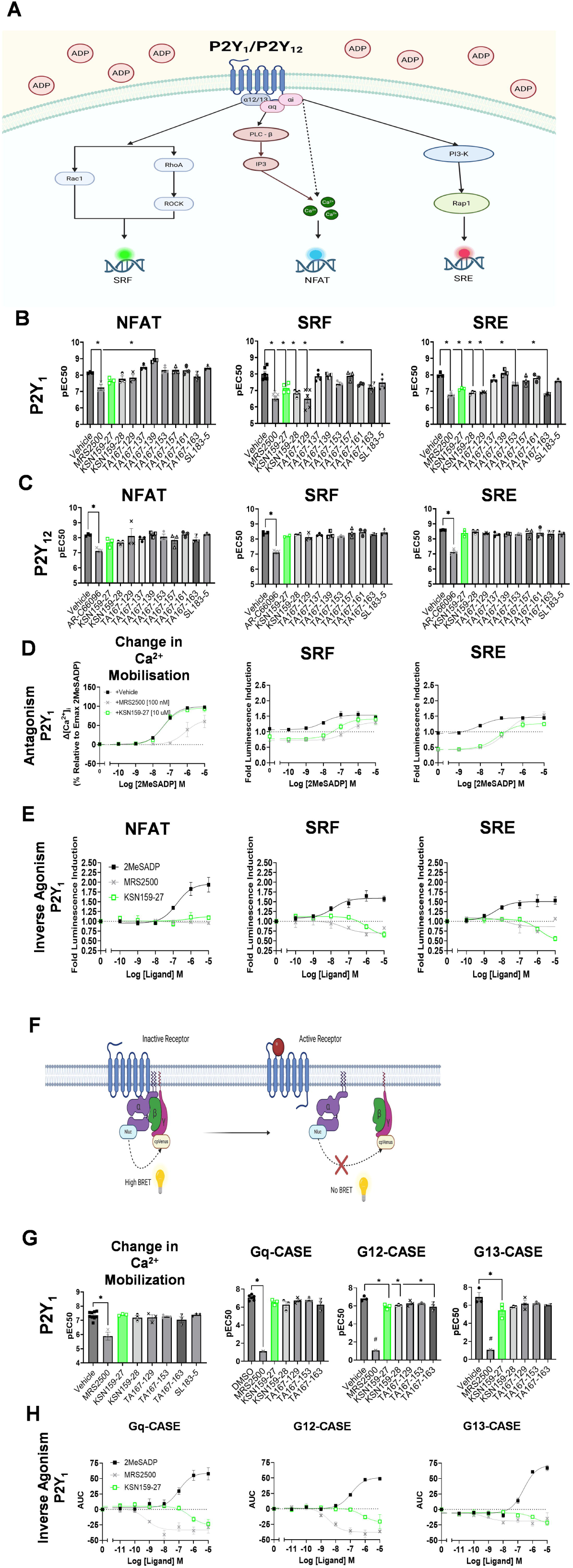
Comparison of antagonistic ability of KSN159-27 and MRS2500 on P2Y_1_R expressing HEK293T cells. Schematic of signalling pathways upstream of transcription factors for genetic reporter assay (**A**). HEK293T cells were transfected to express P2Y_1_R (**B**) or P2Y_12_R (**C**) and experiments were conducted to measure the ability of compounds and MRS2500 or AR-C66096 to antagonise 2MeSADP induced NFAT, SRF-RE or SRE activity. Compounds and MRS2500 or AR-C66096 were used at 10 µM and 1 µM respectively. Analysis of the ability of KSN-159-27 (10 µM) to antagonize 2MeSADP induced Ca^2+^ mobilization, SRE, or SRF-RE activities in comparison to MRS2500 (100 nM) in cells expressing P2Y_1_R (**D**). Investigation of inverse agonism of KSN-159-27 compared to MRS2500 and 2MeSADP in NFAT-RE, SRF, and SRE assays (**E**). (**F**) Schematic of the G-CASE system for G protein activation measurement. Antagonism of KSN-159-27, KSN-159-28, TA-167-129, TA-167-153, and TA-167-163 (10 µM) in comparison to MRS2500 (100 nM) for Ca^2+^ mobilization and G_αq_-, G_α12_-, and G_α13_-CASE activation (**G**). Investigation of inverse agonism of KSN-159-27 compared to MRS2500 and 2MeSADP in G_αq_-, G_α12_-, and G_α13_-CASE activation assays (**H**). Data analysed using two-way ANOVA followed by Dunnett’s test. *P<0.05; n = 3.

Application of 2MeSADP to the NFAT-RE genetic reporter assay displayed activation for both P2Y_1_R and P2Y_12_R expressing cells, evidence of possible G_αq_-PLC activation by P2Y_12_R (although this may not be substantial);^16,31-33^ Therefore, **KSN-159-27**, based on its favourably pathway selective inhibitory characteristics, was chosen for further testing, including antagonism of upstream intracellular Ca^2+^ mobilization. In HEK293T cells endogenously expressing P2Y_1_R, **KSN-159-27** did not antagonize Ca^2+^ mobilization, in contrast to MRS2500 (**Figure 4D**). Due to the endogenous expression of P2Y_1_R, P2Y_12_R could not be tested in isolation.

Since compound **KSN-159-27** consistently displayed selective inhibition of P2Y_1_R-induced signalling activities (**Figure 4B and 4C**), we used the luciferase reporter assays to further illustrate the antagonist relationship potential of **KSN-159-27** compared to MRS2500. Like MRS2500, **KSN-159-27** acted as a competitive antagonist towards SRF and SRE-mediated activities in the presence of the full agonist 2MeSADP, but unlike MRS2500, **KSN-159-27** did not inhibit Ca^2+^ mobilization (**Figure 4D**). As MRS2500 has been reported to be an inverse agonist at the P2Y_1_R,^34^ we used the luciferase reporter assays to probe agonist potential of **KSN-159-27** and show that inverse agonism was evident for SRE and SRF (**Figure 4E**), but not NFAT-RE. In contrast, the incubation of HEK293T cells with the ‘neutral’ P2Y_1_R antagonist MRS2500 displayed a slight inverse agonism to the NFAT and SRF response elements, but not the SRE response element. For comparison, **KSN-159-27** did not reveal any agonist activity to the P2Y_12_ receptor, measuring either G protein dissociation (**Figure S4A**), or genetic reporter responses (**Figure S4B**).The compounds showing significant and selective antagonism of SRF and SRE over NFAT-RE reporter assays at P2Y_1_R (**Figure 4B**), including **KSN-159-27**, **KSN-159-28**, **TA-167-129**, **TA-167-153**, and **TA-167-163**, were also screened for antagonism of P2Y_1_R-dependent Ca^2+^ mobilization. MRS2500 was the only ligand to inhibit the release of intracellular Ca^2+^ verifying that these compounds do not inhibit downstream G_αq_-PLC signalling.

The ability of these compounds to antagonize G protein activation was tested through use of G-protein tri-cistronic activity sensor (G-CASE system) bioluminescence resonance energy transfer (BRET) biosensors, given the greater sensitivity for G_αq_ signalling over other G protein assay systems (**Figure 4F**).^21^ MRS2500 was the only compound to significantly inhibit activation of G_αq_-CASE in cells expressing P2Y_1_R. **KSN-159-27**, **KSN-159-28**, and **TA-167-163** antagonized the activity of G_α12_- and G_α13_-CASE biosensors (**Figure 4G**). MRS2500 was excluded from statistical analysis in the G_α12_- and G_α13_-CASE assays due to ablation of the BRET signal at the concentrations tested. Since the compounds show selective inhibition of P2Y_1_R at the G protein level, the inverse agonist activity of **KSN-159-27** in this assay system was also tested. **KSN-159-27** behaved as an inverse agonist for all G-CASE proteins (**Figure 4H**), including G_αq_-CASE, which is incongruous with data from downstream signalling experiments that show **KSN-159-27** does not antagonize this pathway. This may be due in part to the bioassay system relying on receptor overexpression, where constitutive activity is increased and provides a favourable environment for increased inverse agonism.

Of the two platelet expressed purinergic receptors activated by ADP, **KSN-159-27** displays receptor selectivity for P2Y_1_R compared to P2Y_12_R and competitively and selectively inhibited G_α12/13_ over G_αq/11_ activity induced by 2MeSADP. This occurred because of selective inverse agonism when incubated in the absence of 2MeSADP. **KSN-159-27** can therefore be described as a pathway selective inverse agonist and selective for P2Y_1_R.

Testing effects of the compounds on P2Y_1_R-dependent platelet functions, **KSN-159-27**, **KSN-159-28**, **TA-167-129**, **TA-167-137**, **TA-167-155**, **TA-167-157**, and **TA-157-163** were able to selectively inhibit P2Y_1_R-dependent platelet chemotaxis whilst preserving the ability of platelets to aggregate to ADP (**Table 1**). In general, these data provided evidence of correlations between the inhibition of specific signalling pathways (**Figure 4**), that then dictated their ability to selectively inhibit P2Y_1_R-dependent platelet functions.

**Table 1.** IC_50_ and Imax data provided for synthesized compounds using platelet functional assays. K_B_ (dissociation constant) was used as measure of antagonist affinity to P2Y_1_R.^45,46^ Values calculated via the dose ratio equation A’/A = 1+[Antagonist]/K_B_ for synthesized compounds in Ca^2+^ mobilization, NFAT-RE, SRF, and SRE reporter assays using HEK293T cells of the response induced by 2MeSADP with the presence of the antagonist or DMSO control.

| Compound | Aggregation<br>IC<sub>50</sub> and I<sub>max</sub> (% inhibition) | Ca<sup>2+</sup><br>K<sub>B</sub> | NFAT<br>K<sub>B</sub> | Chemotaxis<br>IC<sub>50</sub> and I<sub>max</sub> (% inhibition) | SRF<br>K<sub>B</sub> | SRE<br>K<sub>B</sub> |
| --- | --- | --- | --- | --- | --- | --- |
| **MRS2500** | **1.13μM** | **3.63nM** | **14.6nM** | **0.17μM, 95-100%** | **3.24nM** | **6.51nM** |
| KSN 159-25 | No effect (1-50μM) | - | - | 44% at 10μM | - | - |
| KSN 159-26 | No effect (1-50μM) | - | - | 50% at 10μM | - | - |
| KSN 159-27 | No effect (1-100μM) | No effect | 5.04μM | 1.24μM, 95-100% | 1.59μM | 1.63μM |
| KSN 159-28 | No effect (1-50μM) | 24.2μM | 7.56μM | 66% at 10μM | 0.73μM | 0.87μM |
| TA 167-129 | No effect (1-50μM) | 25.4μM | 9.91μM | 15.08μM, 80-85% | 0.32μM | 0.94μM |
| TA 167-137 | No effect (1-50μM) | No effect | No effect | 6.23μM, 80-85% | 23.7μM | 11.2μM |
| TA 167-139 | No effect (50μM) | No effect | No effect | No effect (10μM) | 38.2μM | No effect |
| TA 167-153 | No effect (1-50μM) | 50.8μM | No effect | No effect (10μM) | 3.29μM | 3.36μM |
| TA 167-155 | No effect (1-50μM) | 20.6μM | No effect | 2.76μM, 75-80% | 0.27μM | No effect |
| TA 167-157 | No effect (1-50μM) | 18.0μM | No effect | 1.67μM, 75-80% | 30.0μM | 8.31μM |
| TA 167-161 | No effect (50μM) | 78.6μM | No effect | No effect (10μM) | 3.01μM | 17.0μM |
| TA 167-163 | No effect (50μM) | 10.1μM | 13.8μM | 75% at 10μM | 1.67μM | 0.69μM |
| SL 183-5 | No effect (1-50μM) | No effect | No effect | No effect (10μM) | 4.07μM | 7.35μM |
| SL 183-6 | No effect (1-50μM) | - | - | No effect (10μM) | - | - |
| SL 183-7 | No effect (1-50μM) | - | - | No effect (10μM) | - | - |
| SL 183-8 | No effect (1-50μM) | - | - | No effect (10μM) | - | - |

**Table 2.** Overview of antagonist effects of novel compounds on potency at P2Y_1_ signalling pathways.

| Ligand | Gα<sub>i/o</sub>-SRE-RLuc | Gα<sub>12/13</sub>-SRF-GrRLuc | Gα<sub>q</sub>-Ca<sup>2+</sup>-NFAT-Nluc |
| --- | --- | --- | --- |
| MRS2500 | ✓ | ✓ | ✓ |
| KSN159-27 | ✓ | ✓ | X |
| KSN159-28 | ✓ | ✓ | X |
| TA167-129 | ✓ | ✓ | X |
| TA167-137 | X | X | X |
| TA167-139 | X | ↑ | ↑ |
| TA167-153 | ✓ | X | X |
| TA167-157 | X | X | X |
| TA167-161 | X | X | X |
| TA167-163 | ✓ | ✓ | X |
| SL183-5 | X | X | X |

X – did not significantly inhibit potency
✓ - did significantly inhibit potency
↑ - significantly elevated potency

Based on the available 2D interaction maps (**Figure S3**), the active compounds (for example **KMR-82-13**, **KSN-159-27**, **KSN-159-28, TA-167-129**, and **TA-167-163**) generally share an extended binding orientation within the P2Y_1_R orthosteric pocket involving Val194^ECL2^/Arg195^ECL2^/Lys196^ECL2^, Tyr203^45.51^/Thr205^ECL2^/Asp208^ECL2^, Arg287^6.62^/Ala286^6.61^, and Tyr303^7.32^/Tyr306^7.35^/Gln307^7.36^ rather than a single conserved residue interaction (**Figure S5**). This is broadly consistent with a **KMR-82-13/KSN-159-27**-like binding mode. In contrast, inactive compounds (e.g. **TA-167-153**, **TA-167-161**, **SL-183-5**) retain some individual contacts with residues such as Arg195^ECL2^, Lys196^ECL2^, Thr205^ECL2^, Asp208^ECL2^, Arg287^6.62^ or Tyr303^7.32^, but they do not reproduce the same coordinated deeper-pocket binding pattern (**Figure S3 & S5**). Together, these observations suggest that chemotaxis inhibition is unlikely to be driven simply by hydrogen-bond number. Instead, it may depend on pocket depth, ligand orientation, and spatial organisation with a **KMR-82-13**/**KSN-159-27**-like geometry.

### Platelet P2Y_1_R signalling activities and functions are selectively inhibited by KSN-159-27

Incubation of platelets with **KSN-159-27** resulted in a concentration-dependent inhibition of platelet chemotaxis with no effect on ADP-stimulated platelet aggregation in contrast to MRS2500 (**Table 1**, **Figure 5A and 5B**). Considering the difference in antagonist affinity (K_B_) values for these compounds (**Table 1**), a second ‘neutral’ P2Y_1_R antagonist MRS2179 with a lower affinity (K_B_ 109nM) was used.^35,36^ MRS2179 still displayed an ability to inhibit both platelet chemotaxis and aggregation, like MRS2500 (**Figure S6**). Separately, **KSN-159-27** was unable to promote either aggregation (**Figure 5C**) or chemotaxis towards fMLP (**Figure 5D**) in the absence of ADP. This suggests that the inhibitory effect of **KSN-159-27** on ADP-induced chemotaxis, was not due to partial agonism.

**Figure 5.**
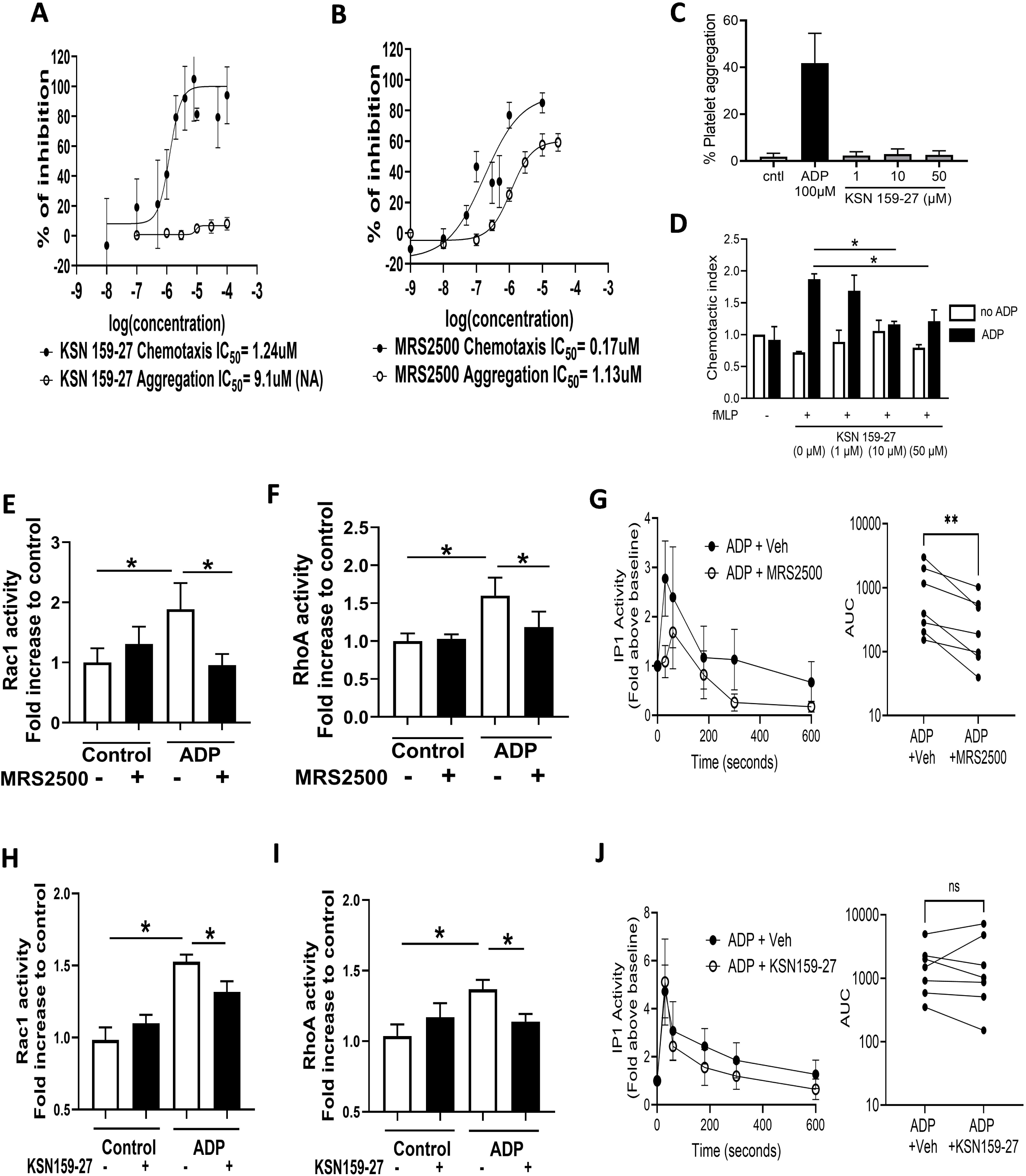
Effect of KSN-159-27 on platelet function *in vitro*. Washed platelets were incubated with KSN159-27 (**A**) or MRS2500 (**B**) and subjected to an inflammatory chemotaxis assay towards fMLP in the presence of ADP (100 nM) (black data points) or hemostatic aggregation assay induced by ADP (10 µM) (white data points). Using these same functional assays, KSN159-27 was incubated with platelets in the absence of ADP, to measure aggregation (**C**) or chemotaxis (**D**). In other assays, platelets were incubated with KSN-159-27 or MRS2500 (10µM) and activated with ADP (10µM). Platelets were then processed for measurement of Rac1 (**E, H**), RhoA (**F, I**) and IP1 activity (**G, J**). Data represented as means +/- SEM, n= 3-9 (**A**,**B**), 3 (**C**,**D**), 5 (**E, H**), 7 (**F, I**), and 8 (**G, J**). Data analysed using two-way ANOVA followed by Dunnett’s test. *P<0.05

Using Up4A as an example of an endogenous nucleotide that can cause P2Y_1_R-dependent platelet chemotaxis but not aggregation,^13^ we show that Rho-GTPase activity increased, and IP1 production was absent (as a measure of upstream PLC activity) to differentiate this effect and compared to ADP (**Figure S7**). To this end, incubation of platelets with **KSN-159-27** significantly inhibited activity of both Rac1 (**Figure 5E,5F**) and RhoA (**Figure 5F,5I**), but in contrast, not IP1 production (**Figure 5G,5J**) induced by ADP. Thus, compared to a ‘neutral’ P2Y_1_R antagonist MRS2500, **KSN-159-27** demonstrated a pathway-specific, functionally selective (biased) inhibition of P2Y_1_R activation of platelets.

### KSN-159-27 inhibits lung inflammation whilst preserving hemostasis *in vivo*

As **KSN-159-27** acts as a pathway-specific inverse agonist at platelet P2Y_1_R, we performed *in vitro* ADME and toxicity analyses in preparation for *in vivo* experimentation. These revealed characteristics best suited to an intravenous route of administration (**Supplementary data: In vitro pharmacokinetic and toxicology assessment**). **KSN-159-27** (0.1-30mg/kg) significantly inhibited total inflammatory cell, neutrophil and monocyte recruitment to the lungs, 4 hours post LPS challenge, similar in magnitude to the inhibition obtained following administration with MRS2500 (3mg/kg) (**Figure 6A and 6B**). Importantly **KSN-159-27** did not affect circulating platelet and neutrophil numbers (**Figure S8).** We next enumerated the tail bleeding time in mice, as a measure of platelet-dependent hemostasis (**Figure 6C**). Whilst the administration of MRS2500 (3mg/kg) led to an expected significant increase in tail bleeding time, no dose of **KSN-159-27**, including 30mg/kg, had an effect (**Figure 6D**). We also demonstrated that the administration of **KSN-159-27**, unlike MRS2500, was unable to suppress the thrombo-embolic response induced by bolus intravenous administration of ADP (**Figure 6E and 6F**). For perspective, the administration of Up4A as an example of an endogenous functionally selective P2Y_1_R agonist was also unable to induce a thromboembolic response when compared to ADP (**Figure S5D**). To determine whether **KSN-159-27** (10mg/kg) might have other non P2Y_1_R mediated effects that could contribute to the inhibition of inflammation, we administered **KSN-159-27** (10mg/kg) into mice selectively deficient in platelet P2Y_1_Rs (C57BL/6J-*P2ry1^em1Kcl^-*Tg(Pf4-icre)Q3Rsko).^12^ However, the repeated administration of **KSN-159-27** (10mins prior, and 6 hours after intranasal administration of LPS) did not further reduce inflammation in these mice (**Figure 6G and 6H**) suggesting that any anti-inflammatory effect of this compound was via direct inhibition of P2Y_1_R- platelet activation.

**Figure 6.**
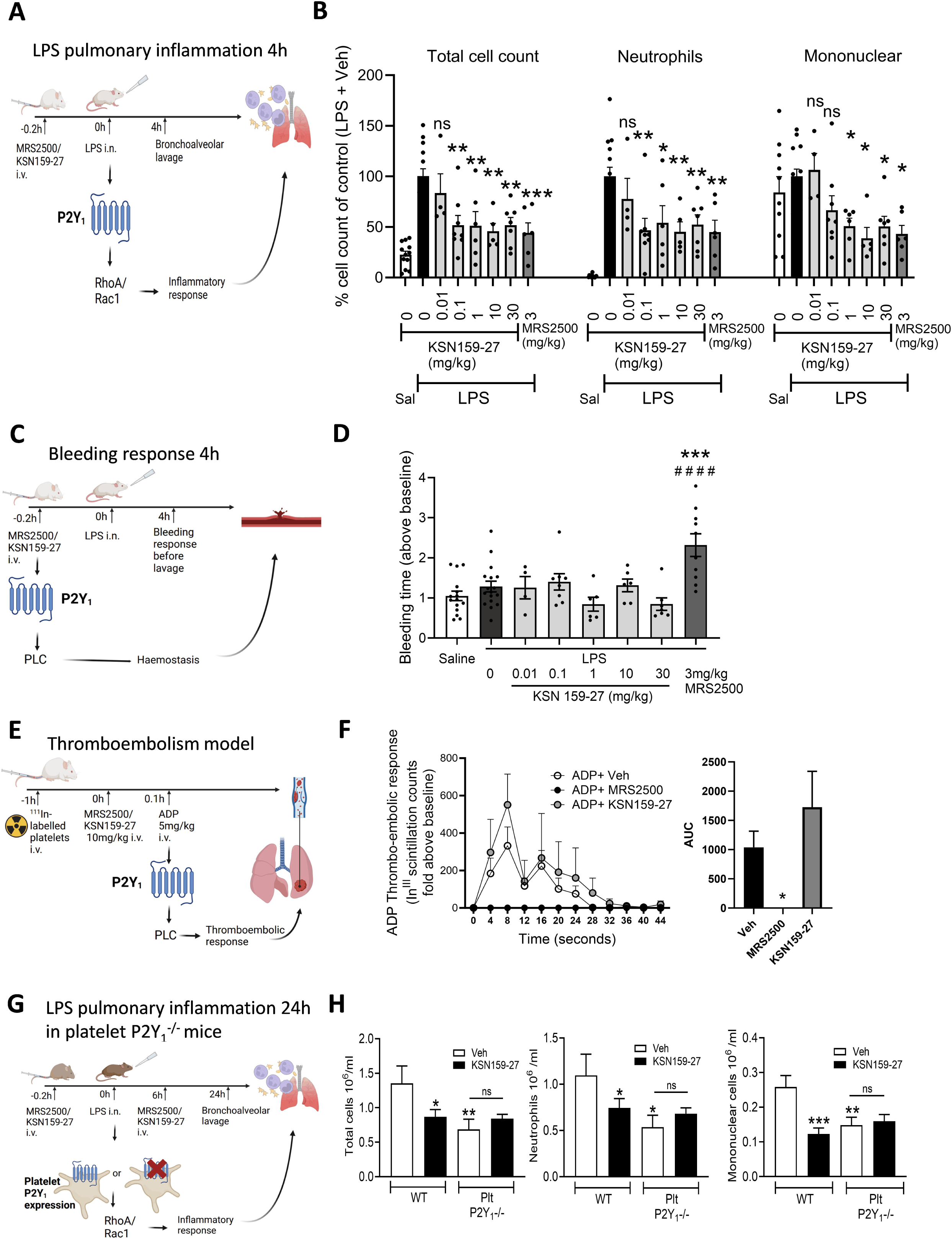
Effect of KSN-159-27 on pulmonary inflammation and hemostasis in mice. Studies were conducted in BALB/C mice to establish whether the intravenous administration of KSN-159-27 (0.01, 0.1, 1, 10, 30 mg/kg) or MRS2500 (3mg/kg) 5-10 minutes before LPS exposure (1mg/kg intranasal) reduced pulmonary inflammatory cell recruitment (**A**). Bronchoalveolar lavages were performed 4 hours after LPS exposure and total cell counts, neutrophils, and mononuclear cells were enumerated (**B**), or tail vein bleeding performed immediately prior to cull in anaesthetised mice (**C, D**). In other experiments, mice were administered ^111^In-labelled platelets intravenously from donor mice (to allow *in vivo* tracking), and then administered KSN-159-27 or MRS2500 (5mg/kg i.v.) before a ADP challenge (5mg/kg i.v.) to induce a pulmonary thromboembolic response in the lung capillary network (**E**), represented as response over time, and area under curve (**F**). Further *in vivo* experiments were conducted with the administration of KSN159-27 (10mg/kg) into mice selectively deficient in platelet P2Y_1_ receptors (C57BL/6J-*P2ry1^em1Kcl^-*Tg(Pf4-icre)Q3Rsko) 10mins prior, and 6 hours after intranasal administration of LPS (1mg/kg intranasal) (**G**). Mice were culled 24 hours later and bronchoalveolar lavages performed for enumeration of total cell counts, neutrophils, and mononuclear cells (**H**). Means +/- SEM n= 4-10 as indicated for groups (**B,D**) because data was amalgamated from 4 separate studies to study dose-response; n= 4-5 (**F**); n= 9 (**H**). *P<0.05, **P<0.01, ***P<0.001 vs LPS group, #### P<0.0001 vs KSN159-27 30mg/kg.

## Discussion

This study reveals a novel tool compound, **KSN-159-27** that exhibits functionally selective, pathway specific inhibition of P2Y_1_R to supress the inflammatory functions of platelets *in vitro* and *in vivo* whilst preserving hemostatic functions, and displays biased inverse agonism at G_12/13_ but not G_αq,_ linked to activation of P2Y_1_R.

Structural modelling of ADP and MRS2500 interactions within the receptor binding pocket has provided important insights into the molecular mechanisms underlying receptor activation and antagonism.^37–40^ All nucleotide ligands share a conserved interaction with Arg195^ECL2^, which destabilizes the ionic lock involving Lys46 and triggers a conformational change in Tyr324^7.53^ that promotes formation of a continuous water channel within the receptor core.^39^ MRS2500 acts as an antagonist by stabilising this channel through insertion into a binding crevice formed between Arg287^6.62^ and Leu44^NTerm^, while forming hydrogen bonds with Asn283^6.58^ located on the boundary of transmembrane helices VI and extracellular loop 3.^37^ Our modelling analyses suggest that biased ligands may disrupt this interaction network in distinct ways. Endogenous nucleotides displaying biased signalling characteristics are predicted to lose interactions with Asn283^6.58^ relative to ADP.^13^ Consistent with this prediction, our nucleotide analogue **KMR-82-13** designed with these biased interactions in mind was also predicted to not directly interact with Asn283^6.58^ or Arg287^6.59^. Loss of this interaction network observed for MRS2500 may therefore explain why **KMR-82-13** fails to suppress platelet hemostatic function.

These observations informed the design of **KSN-159-27**, which notably was predicted to retain only weak van der Waals interactions with Leu44^NTerm^ and Asn283^6.58^, suggesting that partial disengagement from this structural region may be associated with preservation of hemostatic signalling. Interactions shared between endogenous biased nucleotides, **KMR-82-13** and **KSN-159-27** with residues such as Val194^ECL2^, Thr205^ECL2^ and Gln307^7.36^ may influence signalling through G_12/13_-mediated pathways, potentially controlling downstream Rho-GTPase activation and platelet migratory responses. Alternatively, these residues may influence receptor conformations that attenuate coupling to G_αq_-dependent signalling pathways.

Additional differences in receptor interactions further support the existence of distinct ligand-dependent binding modes. For example, MRS2500 uniquely interacts with residues including Cys42^NTerm^, Ala286^6.61^, Gln291^ECL3^ and Asn299^7.28^, whereas **KMR-82-13** and **KSN-159-27** preferentially engage residues such as Val194^ECL2^, Lys196^ECL2^, Thr205^ECL2^ and Gln307^7.36^. The subsequent 200 ns membrane molecular dynamics simulation supported the docking-derived binding model for **KSN-159-27**. The receptor remained structurally stable after equilibration, and **KSN-159-27** remained accommodated within the predicted binding region while undergoing local positional adaptation. Importantly, the ligand retained a stable internal conformation and maintained a recurrent contact network involving residues such as Tyr303^7.32^, Gln291^ECL3^, Arg287^6.62^, Arg195^ECL2^, Thr45^NTerm^, Leu44^NTerm^ and Asp208^ECL2^. Protein RMSF analysis also indicated that the ligand-binding region remained comparatively ordered, while the largest fluctuations were confined mainly to flexible loop and terminal regions outside the principal binding pocket. These MD data therefore support a stable but dynamic binding mode for **KSN-159-27** and suggest that the compound can engage the P2Y_1_R binding pocket without fully reproducing the MRS2500 interaction network associated with broad antagonism.

MRS2500, which displays no functional selectivity for different platelet functions as an antagonist, has been characterized as an inverse agonist, revealing that P2Y_1_R displays basal, constitutive activity.^34^ The significance of this property during hemostasis has yet to be understood. However, the present study demonstrates that achieving pathway-specific inhibition of platelet functions requires ligands capable of inducing biased inverse agonism rather than simple antagonism. These findings highlight the remarkable sensitivity of GPCR signalling outcomes to subtle structural modifications in ligand architecture. Understanding the structure–activity relationships governing biased signalling therefore requires precise characterization of ligand binding poses and receptor interaction networks, as even a single altered amino acid interaction can significantly alter downstream biological responses.^41^ The close alignment of platelet functional data (aggregation vs chemotaxis), and differences in potencies of antagonism for P2Y_1_R-induced NFAT-RE, SRE, SRF-RE, and Ca^2+^ mobilization provides a rationale for future understanding how binding pocket occupancies affect biased ligand activation of this receptor.

The biological basis for this study is that platelet P2Y_1_R via Rho-GTPase signalling is necessary for platelet-leukocyte recruitment and migration as requisite facets of the inflammatory response during host defence.^9,11,12^ Yet, the activation of platelets during inflammation can occur independently of any changes to hemostasis, when platelets aggregate to form thrombi, which in response to ADP is also P2Y_1_R-dependent.^4,42^ Whilst we do not yet understand how platelet activation is controlled during inflammation to preserve hemostasis, we hypothesize that a dynamic purinome that contains biased nucleotides may compete or control signalling pathways operant for hemostasis whilst inducing inflammatory functions. Studies using selective platelet-depletion strategies regularly report a 80-90% suppression of inflammatory events, and 60-80% suppression with selective deficiency/antagonism of platelet P2Y_1_R alone.^9,11,12,43,44^ We have now shown that acute inflammation is inhibited to a similar extent using **KSN-159-27** acting as pathway selective, platelet P2Y_1_R inverse agonist.

To our knowledge, this is the first report of a ‘druggable’ target to selectively supress platelet activation during inflammation whilst preserving hemostasis. Biased inverse agonism of P2Y_1_R represents a novel anti-platelet drug class that is distinct from other anti-platelet drugs such as P2Y_12_R antagonists, aspirin, and integrin αIIbβ3 antagonists used in the clinic prophylactically to reduce thrombosis, but have pronounced haemorrhage liability thereby restricting their potential use in the treatment of inflammatory conditions.

## Author Contributions

D.P., C.M.S., T.A., K.N., K.E., R.T.A., N.G., A.S., E.W., designed and performed the research, analyzed the data, and wrote the paper. R.A. designed research and contributed to project development. K.M.R. conceived and designed research and *in silico* docking analysis and co-wrote the paper. C.P.P., G.L., designed research and co-wrote the paper. S.C.P. conceived the project, designed research and *in vivo* experimentation, wrote the paper.

## Data availability statement

The data that support the findings of this study are available from the corresponding author upon reasonable request. Some data may not be made available because of intellectual property rights, privacy or ethical restrictions.

## Funding Acknowledgement

This research was funded by a Medical Research Council (MRC) project grant [MR/T015845/1] awarded to Prof. Simon Pitchford (PI) and Prof. Miraz Rahman (Co-I); and an endowment awarded to the Pulmonary Pharmacology Unit (Prof. Simon Pitchford and Prof. Clive Page), King’s College London. Prof. Graham Ladds laboratory is funded by the BBSRC (BB/Y513817/1; BB/W014831/1; UKRI1925). C.M.S was funded by a Cambridge Trust International Scholarship in conjunction with the Gonville and Caius Stanley Elmore PhD Studentship. E.W. was funded by an EPSRC studentship co-funded with AstraZeneca (EP/X015785/1, E.W.). We thank Takeda Development Centre Americas Inc. for funding A.S. G.L. is a Royal Society Industry Fellow (INF/R2/212001).

## Conflict of interest

A patent application has been submitted by King’s College London.

## Notes

### Competing Interest Statement

The authors have declared no competing interest.

