## Supplementary data for "Discovery of a pathway-selective platelet P2Y_1_R inverse agonist that suppresses inflammation while preserving hemostasis"

#### Methodology.

##### Molecular docking, molecular dynamics simulation of $P2Y_1R$ and synthesis of KMR-82-13 and non-nucleotide $P2Y_1R$ ligands.

###### Molecular docking studies

Molecular docking was performed using the crystal structure of the human platelet $P2Y_1R$ (P2Y1R; PDB ID: 4XNW). Chain A was prepared in Schrödinger Maestro 2025-4 using the Protein Preparation Wizard.[1] Missing side chains were added, hydrogen-bonding networks were optimised, water molecules outside the binding site were removed, and restrained minimisation was carried out using the OPLS4 force field. Ligands were drawn in ChemDraw and prepared using LigPrep in Schrödinger Maestro 2025-4 (Maestro, Schrödinger, LLC, New York, NY, 2025). Ionisation states were generated at pH 7.0 ± 2.0 using Epik Classic, and ligand geometries were minimised with OPLS4. The receptor grid was generated using Glide Receptor Grid Generation. The docking site was defined using the bound ligand in the receptor structure. The grid centre was set at X = 23.7853, Y = 22.7057 and Z = 15.1414 Å, with an inner box of 10 × 10 × 10 Å and an outer box of 23.2733 × 23.2733 × 23.2733 Å. Flexible ligand docking was then performed using Glide SP with post-docking minimisation. Docking poses were assessed using the Glide docking score and their position within the $P2Y_1R$ binding pocket. The best-scoring and most appropriately positioned poses were selected for comparative interaction analysis.

**Molecular dynamics simulation**

Molecular dynamics simulation of the P2Y1R–KSN-159-27 complex was performed in Schrödinger Maestro/Desmond 2025-2 using the OPLS4 force field.[2] The membrane system was prepared using Desmond System Builder, with automatic placement of the receptor complex in a POPC lipid bilayer. The system was solvated with TIP3P water in an orthorhombic box, using buffer distances of 10 Å in the x and y directions and 15 Å in the z direction. The system was neutralised with 16 $Cl^{-}$ counterions, and 0.15 M NaCl was added. Ion and salt placement was excluded within 5 Å of the protein or ligand.

After Desmond relaxation and equilibration, a 200 ns production simulation was performed under membrane-specific NPγT conditions at 300 K and 1.01325 bar. A 9.0 Å non-bonded cutoff was used, with RESPA time steps of 2/2/6 fs. Trajectory frames and energy data were saved every 100 ps and 10 ps, respectively. RMSD, RMSF, ligand properties, secondary structure and protein–ligand contacts were analysed using the Desmond Simulation Interaction Diagram tools.

**MM-GBSA analysis**

Prime MM-GBSA binding free-energy analysis was performed using frames extracted from the equilibrated part of the 200 ns Desmond trajectory of the $P2Y_1R$–KSN-159-27 complex. The first 20 ns were excluded, and frames were sampled every 2 ns. Calculations were carried out using the VSGB solvation model, OPLS4 force field and implicit membrane treatment. This analysis was used to assess whether KSN-159-27 remained energetically favourable within the $P2Y_1R$ binding pocket during the simulation. Final binding free-energy values will be reported as mean ± SD after completion of the calculation.

#### Chemistry

##### General methods and materials

All reagents and solvents used in the synthesis of the compounds in this report have been obtained from different commercial sources, including Fluorochem, Merck, Alfa Aesar, Fisher Scientific, and Cambridge Bioscience. Thin layer chromatography (TLC) work was performed on silica coated plates (F254 plates, silica gel 60, obtained from Merck). Ultraviolet radiation (UV) was used to visualise the TLC plates at either 254 nm or 365 nm. Flash column chromatography used in the purification of crude mixtures was performed using silica get as normal phase stationary phase (obtained from Merck, 230-400 mesh). Rotary evaporators (obtained from KNF) were utilised for routine evaporation of solvents under vacuum. These rotary evaporators are equipped with RC-600 evaporating system with a SC-920 G vacuum pump. A UN55 oven (Memmert) was used to dry glassware at 200°C.

1H and 13C Nuclear Magnetic Resonance (NMR) experiments were conducted using a 400 MHz spectrometer (obtained from Bruker corp., USA) equipped with a SampleXpress autosampler system. Samples for NMR were prepared using deuterated solvents, while the obtained spectra were analysed using Topspin 7.1 software (Bruker).

**Synthesis of KMR-82-13.**

Commercially available MRS-2500 (1.0 equiv.) was dissolved in 3 mL of anhydrous DMSO under inert gas. Methylpiperazine (1.25 equiv.) and $K_2CO_3$ (6 equiv.) were then added to the solution in a microwave vial, which was sealed with a cap. The reaction mixture was heated at 130 °C and stirred at 600 rpm in a Biotage® Initiator+ Microwave Synthesizer equipped with a pressurized air supply for 15 minutes at 1 bar pressure. After cooling the vial to below 40 °C, the microwave cavity was unlocked, and the reaction mixture was filtered through a 0.45 µm PTFE filter. The crude product was purified using preparative HPLC-MS without further work-up.

**Synthesis of non-nucleotide $P2Y_1R$ ligands**

The synthetic route for the derivatives commences from commercially available starting materials (1 and 2) and progresses through a series of well-defined transformations (Scheme 1). Initially, a cyclization reaction is performed by treating the hydrazine-containing building block and ethyl 3-oxobutanoate with acetic acid and potassium acetate, affording dihydropyrazol-3-one intermediate 3.

Subsequently, intermediate 3 undergoes a nucleophilic aromatic substitution (SnAr) with 1-fluoro-2-nitrobenzene to yield nitro-containing intermediate 5. The nitro group is reduced under two distinct conditions: iron and ammonium chloride afford amine intermediate 6a with the chlorine intact, whereas catalytic hydrogenation produces intermediate 6b with chlorine cleavage.

For Library A, intermediates 6a and 6b are treated with commercially available isocyanate building blocks and triethylamine, forming the urea derivatives as final compounds. For Library B, intermediate 6 is converted to isocyanate 7 using diphosgene and proton sponge, followed by reaction with commercially available amines to yield the final compounds.

This synthetic route offers a modular platform for further derivatization, enabling the preparation of analogues for structure-activity relationship (SAR) studies.

**Synthetic Scheme 1**

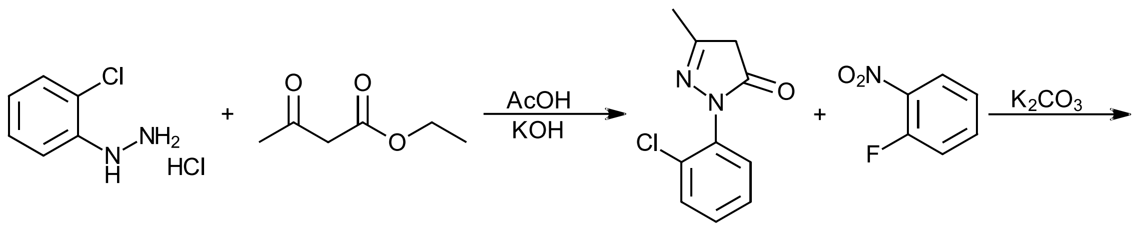

1 2 3 4

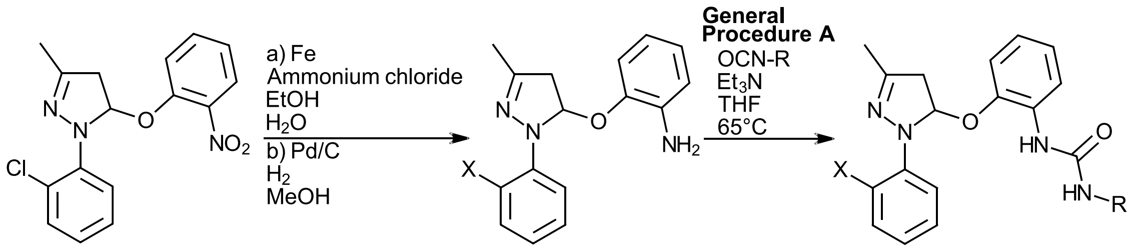

5 6a X = Cl

6b X = H

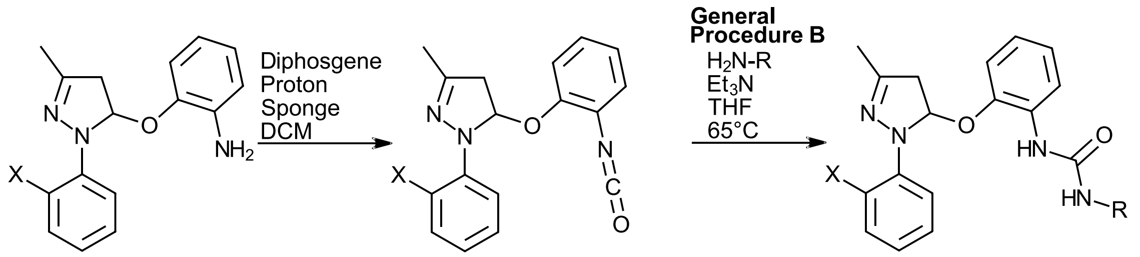

6a X = Cl 7

6b X = H

**Synthetic protocols:**

**Synthesis of 2-(2-chlorophenyl)-5-methyl-2,4-dihydro-3H-pyrazol-3-one (3)**

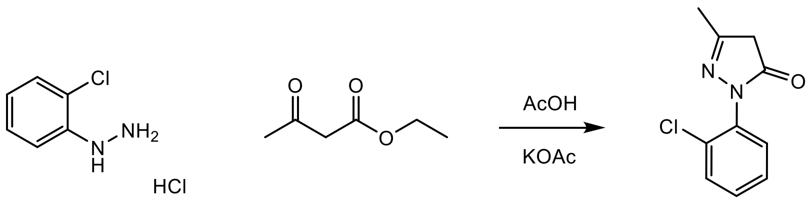

1 2 3

(2-chlorophenyl) hydrazine hydrochloride (4 g, 22.34 mmols, 1.00 eq.) and potassium acetate (3.73 g, 37.98 mmols, 1.70 eq.) were added to a flask. The mixture was dissolved in acetic acid (40 mL) and ethyl acetoacetate (2.91 g, 22.34 mmols, 1.00 eq.) was then added to the flask. The reaction was stirred under reflux and 120°C overnight. When the reaction was finished (monitored by LC-MS), the mixture was cooled down to room temperature, and 500mL of water was added to the mixture. The crude mixture was then extracted using ethyl acetate (150 mL, 3 times) and the combined organic layer was washed with water (200 mL, 2 times) and dried over anhydrous $MgSO_4$. It was then concentrated and purified by column chromatography (80-60% hexane/ethyl acetate) obtaining the product as a yellow solid with 41% yield.

LC-MS: 2.167 min (mass: 209 ($M+H^+$)); $^1$H NMR (400 MHz, $CDCl_3$): δ ppm 7.51-7.49 (1H m), 7.43-7.40 (1H, m), 7.36-7.30 (2H, m), 3.41 (2H, s), 2.19 (3H, s); $^{13}$C NMR (100 MHz, $CDCl_3$): δ 171.44, 156.74, 134.67, 132.04, 130.75, 130.02, 129.05, 127.73, 41.65, 17.31

**Synthesis of 1-(2-chlorophenyl)-3-methyl-5-(2-nitrophenoxy)-1H-pyrazole (5)**

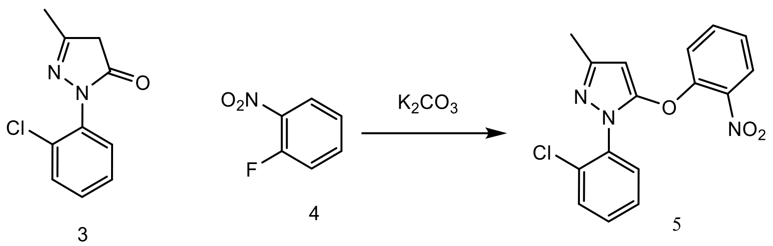

To a solution of 3 (800 mg, 3.83 mmols, 1.00 eq.) in DMF (6 mL) was added potassium carbonate (661.98 mg, 4.79 mmols, 1.25 eq.) and 2-fluoronitrobenzene (0.452 mL, 4.29 mmols, 1.12 eq.). The reaction was stirred at 70°C overnight. The mixture was cooled down to room temperature, and 150mL water was added to the mixture. The crude mixture was then extracted using ethyl acetate (50 mL, 5 times) and the combined organic layer was washed with brine (100 mL, 1 time), dried over anhydrous $MgSO_4$, and concentrated under reduced pressure. The mixture was further purified using column chromatography (100-85% hexane/ethyl acetate) to obtain a brown solid with 21.73% yield.

LC-MS: 3.678 min (mass: 330 ($M+H^+$)); $^1$H NMR (400 MHz, $CDCl_3$): δ ppm 7.88 (1H, $J$ = 8.25 Hz, d), 7.58-7.54 (1H, m), 7.50-7.45 (2H, m), 7.38-7.31 (3H, m), 7.25 (1H, $J$ = 7.77 Hz, t), 5.59 (1H, s), 2.29 (3H, s); $^{13}$C NMR (100 MHz, $CDCl_3$): δ 149.78, 148.98, 134.54, 132.43, 130.64, 130.44, 130.01, 127.70, 126.00, 125.11, 120.69, 90.68, 14.86

**Synthesis of 2-((1-(2-chlorophenyl)-3-methyl-1H-pyrazol-5-yl)oxy)aniline (6a)**

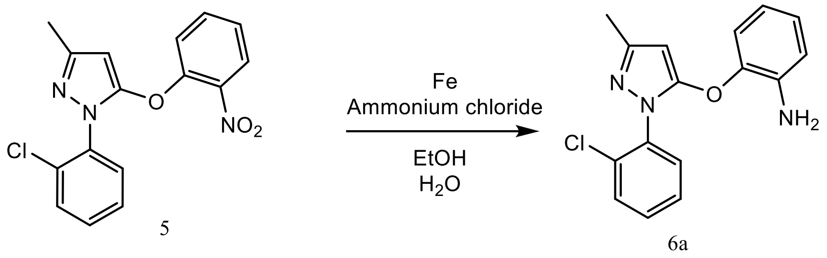

To a solution of 5 (375.19 mg, 1.14 mmols, 1.00 eq.) in ethanol (10.55 mL) and water (3.52 mL), iron (175.8 mg, 3.41 mmols, 3.00 eq.) and ammonium chloride (70.32 mg, 1.37 mmols, 1.2 eq.) were added. The reaction was stirred at 80°C for 2 hours (LC-MS confirmed total consumption of the starting material). The iron was filtered using celite and the mixture was washed with ethyl acetate (total 70 mL, 3 times). The combined organic layer was washed with water (60 mL), dried over anhydrous sodium sulfate, and concentrated under reduced pressure to give a brown solid with 97% yield.

LC-MS: 3.524 min (mass: 300.1 ($M+H^+$)); $^1$H NMR (400 MHz, $CDCl_3$): δ ppm 7.53-7.49 (2H, m), 7.39-7.37 (2H, m), 7.36 (1H, s), 7.08 (1H, *J* = 7.96 Hz, d), 6.97 (1H, *J* = 7.62 Hz, t), 6.76 (1H, *J* = 7.98 Hz, d), 6.70 (1H, *J* = 7.76 Hz, t), 5.44 (1H, s), 3.47 (1H, s), 2.26 (3H, s); $^{13}$C NMR (100 MHz, $CDCl_3$): δ 153.58, 149.85, 143.14, 137.95, 135.82, 132.40, 130.36, 130.04, 127.75, 126.03, 119.45, 118.75, 116.87, 88.72, 14.83.

**Synthesis of 2-((3-methyl-1-phenyl-1H-pyrazol-5-yl)oxy)aniline (6b)**

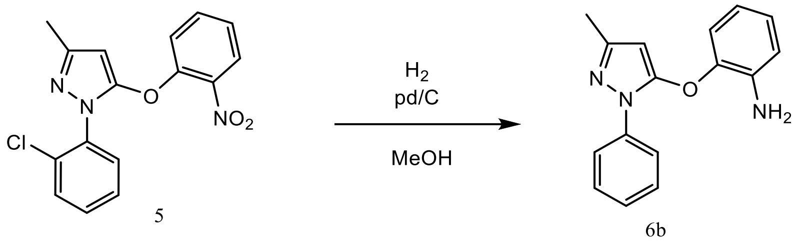

Intermediate 5 (375 mg, 1.13 mmols, 1.00 eq.) was dissolved in methanol (10 mL) and a catalytic amount of Pd/C 10% w/w was added to the reaction mixture in a parr hydrogenator flask and left under pressure of hydrogen gas (around 40 psi) for 3 hours until LC-MS showed completion of reaction. Pd/C of the mixture was removed by filtering on a celite pad, and the organic solvent was evaporated under reduced pressure to obtain 6b as a brown solid with 94% yield.

$^1$H NMR (400 MHz, DMSO-$d_6$) δ 7.61 - 7.70 (m, 2H), 7.44 - 7.56 (m, 3H), 6.87 - 7.00 (m, 3H), 6.74 - 6.79 (m, 1H), 6.50 - 6.58 (m, 1H), 5.39 (s, 1H), 2.14 (s, 3H). $^{13}$C NMR (101 MHz, Acetone) δ 153.5, 148.6, 142.6, 139.2, 131.7, 130.3, 130.1, 130.0, 127.7, 125.7, 119.0, 116.9, 116.2, 88.0, 44.8, 13.7.

**General procedure A for synthesising final compounds:**

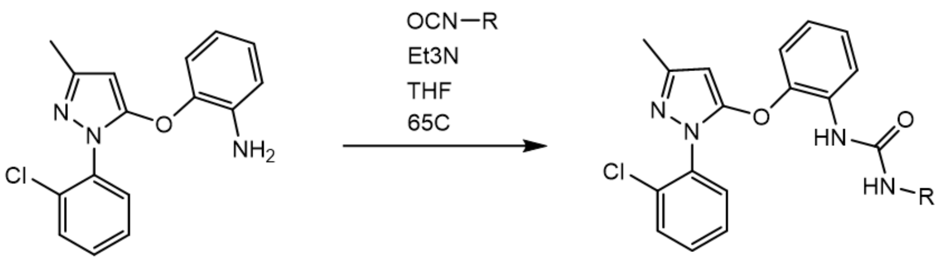

6a

To a solution of 6a (50 mg, 166.8 µmols, 1.00 eq.) in DCM (3 mL), commercially available isocyanate containing building blocks (333.6 µmols, 2.00 eq.) and triethyl amine (0.07 mL, 500.4 µmols, 3.00 eq.) were added. The mixture was stirred at room temperature for 3 hours. When the reaction was finished (confirmed using LC-MS), the solvent was removed under reduced pressure and the resulting residue was purified using column chromatography (100-95% DCM/ethyl acetate) to obtain the final compounds.

**KSN-159-24**

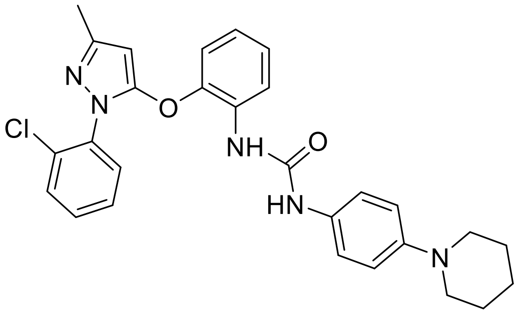

**$^1$H NMR (400 MHz, MeOD)** δ 7.66 – 7.62 (m, 2H), 7.59-7.56 (m, 1H), 7.55 – 7.52 (m, 1H), 7.52-7.51 (m, 1H), 7.50 – 7.44 (m, 1H), 7.28 – 7.24 (m, 1H), 7.06 – 7.03 (m, 1H), 7.02 (dd, *J* = 5.0, 1.4 Hz, 1H), 6.99 (m, 1H), 6.98 – 6.95 (m, 1H), 6.87-6.84 (m, 1H), 6.71-6.69 (m, 1H), 3.67 – 3.63 (m, 2H), 3.50 (t, *J* = 6.6 Hz, 2H), 2.07 (s, 3H), 1.62 – 1.58 (m, 2H), 1.57-1.56 (m, 2H), 1.56 – 1.52 (m, 2H).

**1-(2-((1-(2-chlorophenyl)-3-methyl-1H-pyrazol-5-yl)oxy)phenyl)-3-(4-methoxyphenyl)urea (KSN-159-25):**

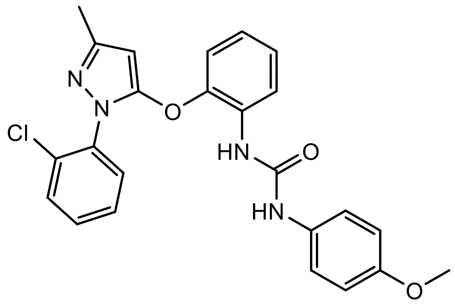

**In a variation to General Procedure A**

To a solution of 2-((1-(2-chlorophenyl)-3-methyl-1H-pyrazol-5-yl)oxy)aniline (50 mg, 166.8 µmol) in THF (5 mL) was added 4-methoxyphenyl isocyanate (49.76 mg, 333.6 µmol) and triethyl amine (50.64 mg, 500.4 µmol). The reaction mixture was stirred at 65 °C for 4 hrs and then cooled to 25 °C. The solvent was removed under reduced pressure and the resulting residue was purified by column chromatography (5-15% ethyl acetate/hexane) to afford 1-(2-((1-(2-chlorophenyl)-3-methyl-1H-pyrazol-5-yl)oxy)phenyl)-3-(4-methoxyphenyl)urea (46 mg, 62%). ESI-MS: 450.03 (M+H)+

**$^{1}$H NMR (400 MHz, MeOD)** δ 8.09 – 8.06 (m, 1H), 7.66-7.64 (m, 2H), 7.48-7.44 (m, 2H), 7.36 – 7.32 (m, 1H), 7.29 – 7.26 (m, 2H), 7.19-7.15 (m, 1H), 7.06 – 7.03 (m, 2H), 6.88 – 6.84 (m, 2H), 3.77 (s, 3H), 2.27 (s, 3H); **$^{13}$C NMR (101 MHz, MeOD)** δ 155.9, 154.2, 151.5, 149.4, 145.8, 131.5, 130.7, 129.8, 129.2, 128.9, 127.7, 127.0, 125.8, 125.0, 124.3, 123.2, 122.5, 121.6, 121.4, 117.4, 116.8, 113.8, 91.5, 54.6, 18.8.

**1-(2-((1-(2-chlorophenyl)-3-methyl-1H-pyrazol-5-yl)oxy)phenyl)-3-(p-tolyl)urea (KSN-159-26):**

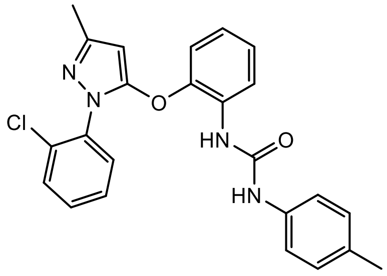

**In a variation to General Procedure A**

To a solution of 2-((1-(2-chlorophenyl)-3-methyl-1H-pyrazol-5-yl)oxy)aniline (50 mg, 166.8 µmol) in THF (5 mL) was added p-tolyl isocyanate (44.42 mg, 333.6 µmol) and triethyl amine (50.64 mg, 500.4 µmol). The reaction mixture was stirred at 65 °C for 4 hrs and then cooled to 25 °C. The solvent was removed under reduced pressure and the resulting residue was purified by column chromatography (5-15% ethyl acetate/hexane) to afford 1-(2-((1-(2-chlorophenyl)-3-methyl-1H-pyrazol-5-yl)oxy)phenyl)-3-(p-tolyl)urea (32 mg, 44%). ESI-MS: 434.72 (M+H)+

**1-(2-((1-(2-chlorophenyl)-3-methyl-1H-pyrazol-5-yl)oxy)phenyl)-3-(naphthalen-2-yl)urea (KSN-159-27):**

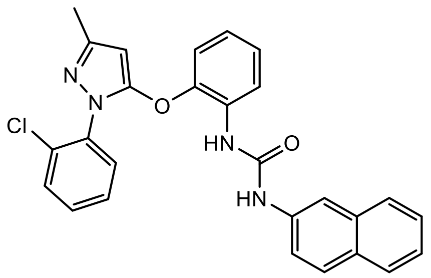

Using General Procedure A obtained 0.051g (reaction yield: 65%) as a white solid.

**$^{1}$H NMR** (400 MHz, DMSO-*d$_6$*): δ ppm 9.03 (1H, s), 8.24 (1H, s), 8.10 (1H, *J* = 8.29 Hz, d), 7.80-7.74 (2H, t), 7.56-7.37 (9H, m), 7.09-7.05 (2H, t), 6.95-6.90 (1H, m), 5.68 (1H, s), 2.14 (3H, s); **$^{13}$C NMR** (100 MHz, DMSO-*d$_6$*): δ 153.27, 152.79, 149.62, 146.08, 138.09, 135.83, 134.68, 131.76, 131.68, 131.15, 130.84, 130.79, 130.12, 129.48, 128.90, 128.42, 127.97, 127.36, 125.86, 125.01, 123.59, 121.58, 120.44, 118.47, 114.38, 91.79, 15.30; **HRMS:** theoretical mass (M+H$^+$) for $C_{27}H_{21}ClN_4O_2$ = 469.1426, observed mass (M+H$^+$) = 469.1425.

**1-(4-butylphenyl)-3-(2-((1-(2-chlorophenyl)-3-methyl-1H-pyrazol-5-yl)oxy)phenyl)urea (KSN-159-28):**

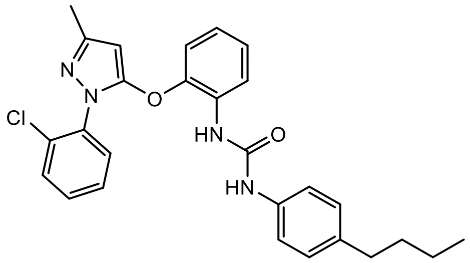

In a variation to General Procedure A

To a solution of 2-((1-(2-chlorophenyl)-3-methyl-1H-pyrazol-5-yl)oxy)aniline (50 mg, 166.8 µmol) in THF (5 mL) was added 4-butylphenyl isocyanate (58.46 mg, 333.6 µmol) and triethyl amine (50.64 mg, 500.4 µmol). The reaction mixture was stirred at 65 °C for 4 hrs and then cooled to 25 °C. The solvent was removed under reduced pressure and the resulting residue was purified by column chromatography (5-15% ethyl acetate/hexane) to afford 1-(4-butylphenyl)-3-(2-((1-(2-chlorophenyl)-3-methyl-1H-pyrazol-5-yl)oxy)phenyl)urea (38 mg, 48%). ESI-MS: 476.21 (M+H)+

**$^{1}$H NMR** (400 MHz, MeOD): δ ppm 8.05-8.02 (1H, m), 7.63-7.61 (2H, m), 7.45-7.41 (2H, m), 7.33-7.30 (1H, m), 7.26-7.24 (2H, m), 7.17-7.14 (1H, m), 7.14-7.12 (2H, m), 7.08-7.06 (2H, m), 2.52 (2H, *J* = 7.2 Hz, t) 2.24 (3H, s), 1.58-1.50 (2H, m), 1.35-1.28 (2H, m), .92-.89 (3H, m);

**1-(2-((1-(2-chlorophenyl)-3-methyl-1H-pyrazol-5-yl)oxy)phenyl)-3-(3,5-dimethoxyphenyl)urea (SL-183-5)**

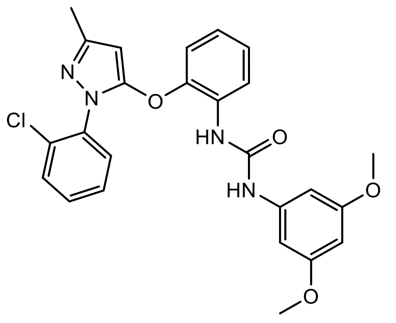

**Using General Procedure A obtained 0.048g (reaction yield: 60%) as a white solid.**

**$^1$H NMR** (400 MHz, DMSO-$d_6$): δ ppm 9.14 (1H, s), 8.19 (1H, s), 8.07 (1H, *J* = 8.46 Hz, d), 7.62-7.58 (2H, m), 7.49-7.42 (2H, m), 7.12-7.08 (2H, t), 6.99-6.95 (1H, m), 6.63-6.27 (2H, d), 6.14 (1H, s), 5.7 (1H, s), 3.71 (6H, s), 2.19 (3H, s); **$^{13}$C NMR** (100 MHz, DMSO-$d_6$): δ 161.64 (2C), 153.02, 152.81, 149.64, 146.06, 142.18, 135.84, 131.76, 131.69, 131.17, 130.79 (2C), 128.92, 125.85, 123.58, 121.59, 118.46, 97.34 (2C), 95.15, 91.77, 56.02 (2C), 15.31; **HRMS:** theoretical mass (M+H$^+$) for $C_{25}H_{23}ClN_4O_4$ = 479.1481, observed mass (M+H$^+$) = 479.1462.

**1-(2-((1-(2-chlorophenyl)-3-methyl-1H-pyrazol-5-yl)oxy)phenyl)-3-(4-cyanophenyl)urea (SL-183-6)**

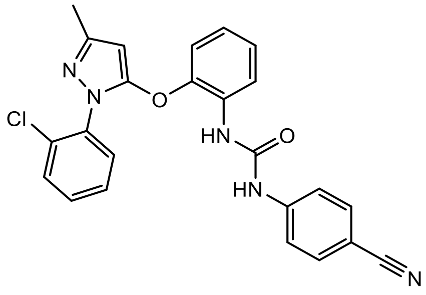

Using General Procedure A obtained 0.042g (reaction yield: 57%) as a brown solid.

**$^1$H NMR** (400 MHz, DMSO-$d_6$): δ ppm 9.63 (1H, s), 8.39 (1H, s), 8.02 (1H, *J* = 8.64 Hz, d), 7.73-7.71 (2H, q), 7.60-7.57 (4H, m), 7.14-7.10 (2H, t), 7.01 (1H, *J* = 7.86 Hz, t), 5.73 (1H, s), 2.19 (3H, s); **$^{13}$C NMR** (100 MHz, DMSO-$d_6$): δ 152.79, 152.65, 149.57, 146.44, 144.95, 135.79, 134.28 (2C), 131.76, 131.63, 131.09, 130.82, 130.26, 128.84, 125.81, 124.17, 121.99, 120.22, 118.93 (2C), 118.45, 104.38, 91.84, 15.29; **HRMS:** theoretical mass (M+H$^+$) for $C_{24}H_{18}ClN_5O_2$ = 444.1222, observed mass (M+H$^+$) = 444.1223.

**1-2-((1-(2-chlorophenyl)-3-methyl-1H-pyrazol-5-yl)oxy)phenyl)-3-(2,3-dihydro-1H-inden-5-yl)urea (SL-183-7)**

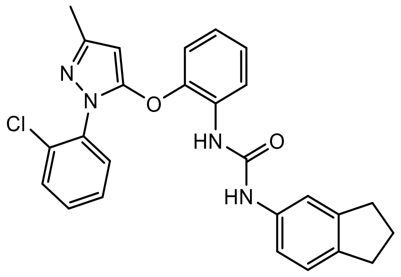

Using General Procedure A obtained 0.019g (reaction yield: 25%) as a white solid.

**$^{1}$H NMR** (400 MHz, methonal-*$d_4$*): δ ppm 8.01 (1H, *J* = 7.98 Hz, d), 7.59-7.41 (4H, m), 7.31 (1H, s), 7.18-7.03 (5H, m), 5.72 (1H, s), 2.93-2.87 (4H, q), 2.29 (3H, s), 2.14-2.07 (2H, q); **$^{13}$C NMR** (100 MHz, methonal-*$d_4$*): δ 154.24, 151.23, 147.26, 146.06, 140.08, 138.30, 136.16, 133.44, 132.06, 131.42, 131.22, 131.04, 128.87, 126.28, 125.36, 124.41, 123.02, 119.00 (2C), 117.02, 91.91, 33.95, 33.16, 26.74, 14.26; **HRMS:** theoretical mass (M+H$^+$) for $C_{26}H_{23}ClN_4O_2$ = 459.1582, observed mass (M+H$^+$) = 459.1567.

**1-(2-((1-(2-chlorophenyl)-3-methyl-1H-pyrazol-5-yl)oxy)phenyl)-3-(3,4-difluorophenyl)urea (SL-183-8)**

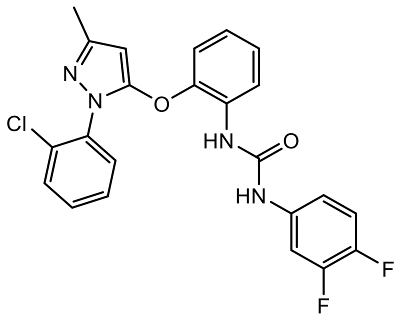

Using General Procedure A obtained 0.066g (reaction yield: 87%) as a brown solid.

**$^{1}$H NMR** (400 MHz, methonal-*$d_4$*): δ ppm 7.86 (1H, *J* = 8.01 Hz, d), 7.70 (1H, *J* = 8.01 Hz, d), 7.63 (1H, *J* = 8.01 Hz, d), 7.57-7.53 (2H, m), 7.47 (1H, *J* = 7.51 Hz, t), 7.26-7.21 (2H, m), 7.18-7.13 (2H, t), 7.08-7.02 (1H, m), 5.86 (1H, s), 2.35 (3H, s); **$^{13}$C NMR** (100 MHz, methonal-*$d_4$*): δ 155.51, 154.67, 151.05, 147.34, 134.02, 133.54, 133.24, 131.68, 131.38, 131.03, 129.28, 127.25, 125.52, 124.30, 119.89, 87.50, 13.44; **HRMS:** theoretical mass (M+H$^+$) for $C_{23}H_{17}ClF_2N_4O_2$ = 455.1081, observed mass (M+H$^+$) = 455.1071.

**General procedure B for synthesising final compounds (one pot synthesis):**

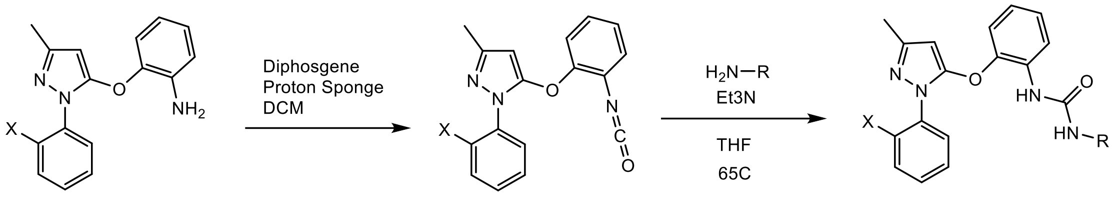

6

7

To a solution of 6 (0.234 mmols, 1.00 eq.) in DCM (5 mL), diphosgene (0.178 mmols, 0.76 eq.) was added at 0 °C. Proton sponge (0.468 mmols, 2 eq.) was separately dissolved in DCM (2 mL) and added to the reaction mixture at 0 °C. The mixture was stirred while allowing the temperature to gradually increase to room temperature. After confirming the formation of the isocyanate 7 using LC-MS, commercially available aryl amines (0.304 mmols, 1.3 eq.) and triethyl amine (0.351mmols, 1.5 eq.) were added. The mixture was stirred at 65 °C for 3 hours. The reaction was then quenched with water and extracted with ethyl acetate (30 ml, 3 times). The combined organic layer was then washed with brine (30mL, once), dried over sodium sulfate, and then concentrated under reduced pressure. The resulting residue was purified using column chromatography (100-70% DCM/ethyl acetate) to obtain the final compounds.

**1-(2-((1-(2-chlorophenyl)-3-methyl-1H-pyrazol-5-yl)oxy)phenyl)-3-(1-methyl-1H-benzo[d]imidazol-5-yl)urea (TA-167-129)**

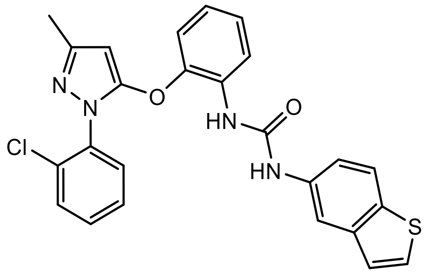

Obtained 0.03g (reaction yield: 27%) as a white solid.

**$^{1}$H NMR** (400 MHz, DMSO-d6) δ 9.25 (s, 1H), 8.25 (s, 1H), 8.12 (d, J = 7.79 Hz, 1H), 8.08 (d, J = 1.83 Hz, 1H), 7.89 (d, J = 8.71 Hz, 1H), 7.73 (d, J = 5.32 Hz, 1H), 7.59 - 7.65 (m, 2H), 7.44 - 7.50 (m, 2H), 7.40 (d, J = 5.50 Hz, 1H), 7.32 (dd, J = 1.93, 8.71 Hz, 1H), 7.11 (d, J = 7.79 Hz, 2H), 6.95 - 7.01 (m, 1H), 5.73 (s, 1H), 2.20 (s, 3H). **$^{13}$C NMR** (101 MHz, DMSO-d6) δ 152.8, 152.3, 149.1, 145.5, 140.6, 136.9, 135.3, 133.3, 131.3, 131.2, 130.7, 130.5, 130.3, 128.6, 128.4, 125.3, 124.4, 123.2, 122.9, 121.0, 118.0, 117.2, 112.7, 91.3, 14.8. **HRMS:** theoretical mass (M+H$^{+}$) for $C_{25}H_{19}ClN_4O_2S$= 475.0984, observed mass (M+H$^{+}$) = 475.0990.

**1-(2-((1-(2-chlorophenyl)-3-methyl-1H-pyrazol-5-yl)oxy)phenyl)-3-(quinazolin-7-yl)urea (TA-167-137)**

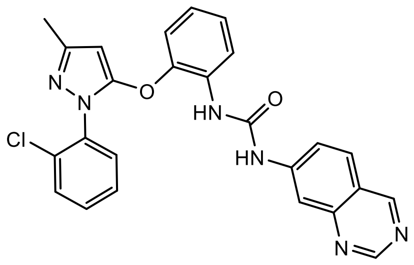

Using General Procedure B obtained 0.021g (reaction yield: 19%) as a brown solid.

**$^1$H NMR** (400 MHz, DMSO-d6) d 9.80 (s, 1H), 9.36 - 9.42 (m, 1H), 9.16 (d, J = 2.38 Hz, 1H), 8.48 (s, 1H), 8.23 (s, 1H), 8.01 - 8.13 (m, 2H), 7.55 - 7.65 (m, 3H), 7.40 - 7.50 (m, 2H), 7.11 - 7.21 (m, 2H), 6.99 - 7.09 (m, 1H), 5.72 - 5.80 (m, 1H), 2.20 (d, J = 2.20 Hz, 3H). **$^{13}$C NMR** (101 MHz, DMSO) δ 157.95, 156.07, 155.80, 154.81, 152.42, 152.40, 152.17, 149.13, 146.01, 145.26, 131.27, 130.63, 129.74, 129.27, 128.38, 125.38, 123.79, 121.58, 120.14, 118.34, 118.00, 112.17, 103.85, 91.44, 14.80.**HRMS:** theoretical mass (M+H$^+$) for $C_{25}H_{19}ClN_6O_2$ = 471.1325, observed mass (M+H$^+$) = 471.1331

**TA-167-139**

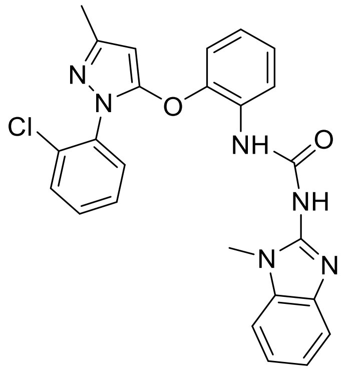

Using general procedure B, obtained 0.105 g (reaction yield: 37%) as a white solid.

**$^1$H NMR (400 MHz, DMSO)** δ 12.04 (d, *J* = 21.4 Hz, 1H), 8.43 – 8.27 (m, 1H), 7.71 (dd, *J* = 7.5, 2.0 Hz, 1H), 7.66 – 7.60 (m, 1H), 7.59 – 7.48 (m, 2H), 7.44 – 7.38 (m, 1H), 7.38 – 7.29 (m, 1H), 7.20-7.17 (m, 2H), 7.17 – 7.12 (m, 2H), 7.02 – 6.94 (m, 1H), 3.72 (s, 1H), 3.52 (s, 2H), 2.18 (d, *J* = 13.0 Hz, 3H).

**1-(benzo[d]thiazol-5-yl)-3-(2-((1-(2-chlorophenyl)-3-methyl-1H-pyrazol-5-yl)oxy)phenyl)urea (TA-167-153)**

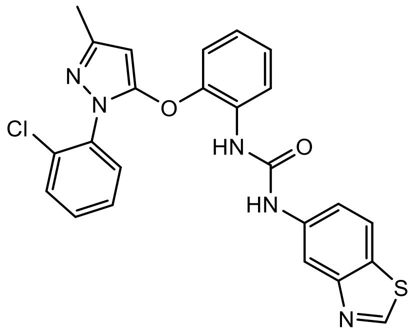

Using General Procedure B obtained 0.033g (reaction yield: 30%) as a brown solid.

**$^{1}$H NMR** (400 MHz, DMSO-$d_6$) d 9.39 (s, 1H), 9.36 (s, 1H), 8.27 - 8.34 (m, 2H), 8.11 (d, *J* = 8.34 Hz, 1H), 8.04 (d, *J* = 8.71 Hz, 1H), 7.58 - 7.65 (m, 2H), 7.43 - 7.49 (m, 2H), 7.41 (dd, *J* = 2.02, 8.71 Hz, 1H), 7.12 (d, *J* = 7.89 Hz, 2H), 6.96 - 7.04 (m, 1H), 5.74 (s, 1H), 2.20 (s, 3H), 1.99 (s, 1H). **$^{13}$C NMR** (101 MHz, DMSO) δ 157.36, 154.23, 152.81, 152.31, 149.22, 145.71, 138.60, 135.25, 131.26, 131.22, 130.61, 130.29, 130.16, 128.41, 127.22, 125.41, 123.34, 122.91, 121.39, 117.99, 111.99, 91.25, 14.74. **HRMS:** theoretical mass (M+H$^+$) for $C_{24}H_{18}ClN_5O_2S$ = 476.0937, observed mass (M+H$^+$) = 476.0943.

**1-(2-((1-(2-chlorophenyl)-3-methyl-1H-pyrazol-5-yl)oxy)phenyl)-3-(1-methyl-1H-indol-5-yl)urea (TA-167-155)**

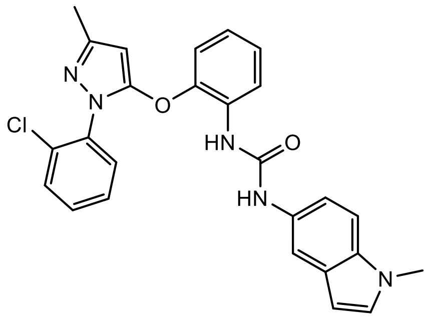

Using General Procedure B obtained  0.027g (reaction yield: 25%) as a brown solid.

**$^{1}$H NMR** (400 MHz, Acetone) δ 8.35 – 8.28 (m, 1H), 8.16 (s, 1H), 7.75 (d, J = 2.0 Hz, 1H), 7.69 (s, 1H), 7.56 (dd, J = 6.8, 1.5 Hz, 1H), 7.43 (d, J = 6.3 Hz, 2H), 7.31 (d, J = 8.6 Hz, 1H), 7.23 – 7.16 (m, 2H), 7.16 – 7.07 (m, 2H), 6.96 (dd, J = 8.6, 7.0 Hz, 1H), 6.38 (d, J = 3.7 Hz, 1H), 5.66 (s, 1H), 3.82 (d, J = 1.5 Hz, 3H), 2.23 (s, 3H). **$^{13}$C NMR** (101 MHz, Acetone) d 157.4, 148.9, 145.0, 142.5, 139.6, 135.6, 130.3, 130.1, 129.7, 128.8, 127.6, 124.8, 121.9, 120.4, 119.3, 117.3, 112.4, 109.3, 100.3, 94.6, 90.4, 89.0, 57.6, 53.9, 32.0, 13.8. **HRMS:** theoretical mass (M+H$^+$) for $C_{26}H_{22}ClN_5O_2$ = 472.1531, observed mass (M+H$^+$) = 472.1535

**1-(1H-benzo[d]imidazol-5-yl)-3-(2-((1-(2-chlorophenyl)-3-methyl-1H-pyrazol-5-yl)oxy)phenyl)urea (TA-167-157)**

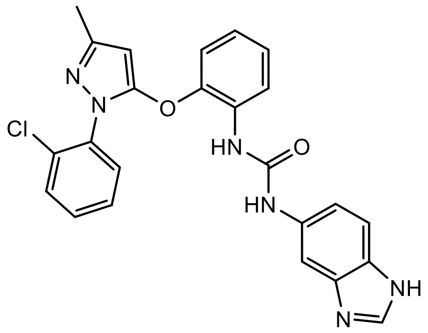

Using General Procedure B obtained 0.024g (reaction yield: 22%) as a white solid.

**$^1$H NMR** (400 MHz, DMSO-d$_6$) d 8.11 (s, 1H), 8.06 (dd, *J* = 1.42, 8.12 Hz, 1H), 7.85 (d, *J* = 1.74 Hz, 1H), 7.57 - 7.62 (m, 2H), 7.50 (d, *J* = 8.62 Hz, 1H), 7.42 - 7.47 (m, 2H), 7.06 - 7.12 (m, 2H), 7.01 - 7.05 (m, 1H), 6.93 - 7.01 (m, 1H), 5.69 (s, 1H), 2.72 (s, 2H), 2.15 - 2.21 (m, 3H). **$^{13}$C NMR** (101 MHz, DMSO) δ 155.96, 152.45, 151.15, 149.27, 146.08, 145.29, 135.20, 131.24, 130.60, 130.27, 129.58, 129.27, 128.40, 125.45, 124.02, 121.82, 121.45, 121.09, 118.02, 112.08, 91.38, 14.72. **HRMS:** theoretical mass (M+H$^+$) for $C_{24}H_{19}ClN_6O_2$ = 459.1323, observed mass (M+H$^+$) = 459.1331.

**1-(2-((3-methyl-1-phenyl-1H-pyrazol-5-yl)oxy)phenyl)-3-(1-methyl-1H-benzo[d]imidazol-5-yl)urea (TA-167-161)**

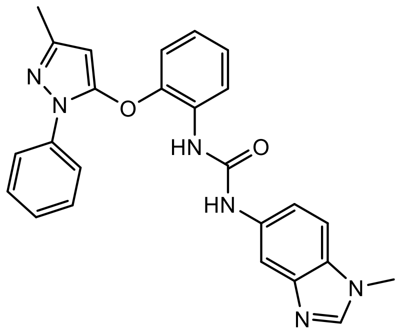

Using General Procedure B obtained 0.063g (reaction yield: 61%) as a brown solid.

**$^1$H NMR** (400 MHz, Acetone) d 8.55 (br. s., 1H), 8.46 (d, *J* = 8.07 Hz, 1H), 8.12 (d, *J* = 11.19 Hz, 1H), 8.00 (s, 1H), 7.93 (s, 1H), 7.79 (d, *J* = 8.16 Hz, 2H), 7.47 (t, *J* = 7.75 Hz, 2H), 7.34 - 7.43 (m, 2H), 7.26 - 7.31 (m, 1H), 7.20 (t, *J* = 7.84 Hz, 1H), 7.09 (d, *J* = 8.16 Hz, 1H), 7.00 (t, *J* = 7.70 Hz, 1H), 5.62 (s, 1H), 3.89 (s, 3H), 2.54 (s, 2H), 2.23 (s, 3H). **$^{13}$C NMR** (101 MHz, Acetone) d 155.8, 148.5, 144.9, 144.5, 141.1, 135.2, 135.1, 129.0, 128.1, 126.2, 125.1, 125.2, 122.2, 121.4, 120.5, 120.4, 117.4, 115.5, 114.4, 109.5, 96.5, 91.8, 73.4, 30.3, 15.3. **HRMS:** theoretical mass (M+H$^+$) for $C_{25}H_{22}N_6O_2$ = 439.1869, observed mass (M+H$^+$) = 439.1877.

**1-(2-((1-(2-chlorophenyl)-3-methyl-1H-pyrazol-5-yl)oxy)phenyl)-3-phenylurea (TA-167-163)**

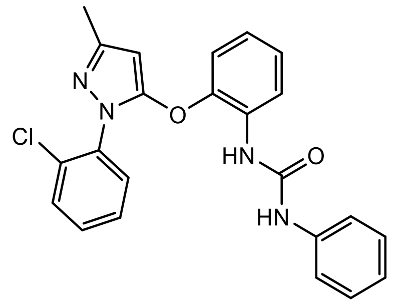

Using General Procedure B obtained 0.031g (reaction yield: 32%) as a yellow solid.

**$^1$H NMR** (400 MHz, Acetone) d 8.23 (br. s., 1H), 8.11 (d, *J* = 8.16 Hz, 1H), 7.69 (s, 1H), 7.38 - 7.46 (m, 2H), 7.36 (d, *J* = 9.26 Hz, 3H), 7.29 - 7.33 (m, 2H), 7.14 (t, *J* = 7.75 Hz, 2H), 6.93 - 7.03 (m, 2H), 6.85 (dt, *J* = 3.76, 7.43 Hz, 2H), 5.55 (s, 1H), 2.69 (s, 3H), 2.65 (s, 1H), 2.08 (s, 3H), 1.87 - 1.95 (m, 10H). **$^{13}$C NMR** (101 MHz, Acetone) d 152.0, 148.9, 145.3, 144.2, 139.8, 135.6, 134.3, 131.5, 130.4, 130.1, 129.8, 128.7, 127.7, 124.7, 122.3, 122.2, 120.6, 120.5, 118.5, 118.4, 117.2, 90.7, 13.8. **HRMS:** theoretical mass ($M+H^+$) for $C_{23}H_{19}ClN_4O_2$ = 419.1262, observed mass ($M+H^+$) = 419.1269.

### Biology

**Materials**

Acid Citrate Dextrose (ACD)-A Vacuette tubes (Cat #455055) were purchased from Greiner Bio-One. ADP (Cat #01905), N-formylmethionyl-leucyl-phenylalanine (f-MLP) (Cat #F3506) and prostaglandin $E_1$ ($PGE_1$) were all purchased from Sigma Aldrich. The HTS Transwell 96-well plates (3µm pore size) (Cat #10077792) and RPMI 1640 cell media with L-glutamine (Cat #12004997) were purchased from Fisher Scientific. The $P2Y_1$ antagonist, MRS2500 (Cat #2159/1) was purchased from Bio-Techne. The $P2Y_{12}$ antagonist, AR-C66096 (Cat #3321) was purchased from Tocris. HEK293T cells were grown in Dulbecco's Modified Eagle Medium/Hams F12 Nutrient Mix with Glutamax® (DMEM/F12) or DMEM/F-12 media without phenol red purchased from Gibco™ (Thermo Fisher Scientific; UK). Foetal Bovine Serum (FBS) was purchased from several manufacturers due to fluctuations in price and availability. Antibiotic-antimycotic (AA) solution was purchased at 100X from Sigma-Aldrich (Merck; Dorset, UK). Trypsin-EDTA solution was purchased at 10X concentration from Sigma-Aldrich (Merck, Dorset, UK) and diluted to 1X with phosphate buffered saline (PBS) for a final concentration of 0.05%. Poly-L-lysine (PLL) was purchased from ScienCell (California, USA). Polyethyleneimine (PEI) 25kDA was purchased from Polysciences Inc. (Germany).

**Human platelet isolation**

For all studies, blood was collected in accordance with local ethical approval from King's College London (Research Ethics Committee Reference: 10/H0807/99) and adhered to regulations outlined by the Human Tissue Act 2004 as previously described.[3,4] Blood was collected from an antecubital vein using ACD-A Vacuette tubes from healthy male and female volunteers who had not taken non-steroidal anti-inflammatory drugs (NSAIDs) or other anti-inflammatory drugs in the previous seven days, and who were not prescribed anti-platelet drugs. Whole blood was centrifuged at 133 g for 20 minutes at room temperature. An aliquot of the upper platelet-rich plasma (PRP) layer was then used for aggregation studies *(see below)*. To the remaining PRP, 2.5 µM Prostaglandin $E_1$ ($PGE_1$) was added before centrifuging at 800 g for 10 minutes at room temperature. Platelet-poor plasma (PPP) was removed and used for aggregation studies *(see below)*. The platelet pellet was resuspended in RPMI 1640 media, again adding 2.5 µM $PGE_1$ and centrifuging at 800 g for 10 minutes at room temperature. Platelets were then adjusted to a final concentration of $5x10^7$ platelets/ml in RPMI 1640 media using an Improved Neubauer chamber (Hawksley & Sons Ltd) for chemotaxis studies *(see below)*.

##### ***In vitro*** **platelet aggregation**

The various experimental compounds (Table 1) were investigated for their effects on platelet aggregation in response to ADP (100 µM) quantified by light transmission aggregometry of stimulated PRP at 595nm at 37 ºC using a SpectraMax 340PC shaking plate reader (Molecular Devices) as previously described.[3,4] Briefly, PRP was stimulated with vehicle (phosphate buffered saline- PBS) or individual agonists and immediately loaded onto the plate reader. Vehicle stimulated PPP was also used as a control. Measurements were taken at 15-second intervals for 6 minutes under shaking conditions.

##### ***In vitro*** **platelet chemotaxis**

Inflammatory platelet function downstream of purinergic receptor activation induced by ADP (100 nM) was also investigated through *in vitro* platelet chemotaxis, as previously described, but with minor amendments.[3,4] Washed platelets ($5x10^7$/ml) were treated with 2 mM $CaCl_2$ before stimulation with vehicle (PBS) or ADP for 5 minutes at room temperature. In some studies, platelets were incubated with antagonists (Table 1) for 10 minutes at room temperature, prior to stimulation. Platelets (80µl) were then added to the top insert of the 96-well transwell plate, with chemoattractant in the bottom well (0/30nM fMLP in RPMI 1640 cell media). Following 90 minutes incubation at 37 ºC, media from the bottom chamber was stained with Stromatol (1:0.5) and platelets were quantified using an

Improved Neubauer haemocytometer and a Leica DM 2000 LED microscope with an x40 objective lens.

**RhoA, Rac1 activity assays**

Platelets ($2.5x10^9$ cells/ml), were resuspended in Tyrode's buffer with $Ca^{2+}$ (2 mM) and were stimulated with ADP or Up4A (10 µM), and in some conditions, pre-incubated with $P2Y_1$ antagonist MRS2500 or KSN-159-27. After ADP stimulation (30 seconds for Rac1 measurement and 1 minute for RhoA measurement), cells were lysed with an equal volume of ice-cold lysis buffer, following the manufacturer's protocol. Lysates were vortexed at 4°C, centrifuged at 12,000g for 3 minutes at 2°C to remove debris, and then applied to a 96-well Rac1 or RhoA-GTP binding plate (Kits from Cytoskeleton Inc BK124, and BK128 respectively). Plates were incubated at 4°C on a shaker at 400 RPM for 30 minutes, washed twice with wash buffer, and then incubated with antigen presenting buffer for 2 minutes before being washed again. Primary antibodies for Rac1 or RhoA were added and incubated for 45 minutes at room temperature. After washing three times, secondary antibodies were added and incubated for 45 minutes at room temperature. Following additional washes, HRP detection agents were added, and the plate was incubated at room temperature for 10 minutes (Rac1) or at 37°C for 20 minutes (RhoA). The reaction was stopped, and absorbance was measured at 490 nm.

**IP1 production assay**

Intracellular concentrations of IP-1 were quantified in platelets using the HTRF-IP1 Gq detection kit (62IPAPEB, Revvity Inc) following the manufacturer's instructions. Briefly, isolated platelets were resuspended in stimulation buffer containing 2 mM $Ca^{2+}$ and transferred to an HTRF 96-well low volume plate (white) at a density of $1.75x10^7$ cells per well in 7 µL of stimulation buffer. Platelets were activated with ADP or Up4A (10 µM) for 30 seconds, 60 seconds, 3 minutes, 5 minutes, 10 minutes, and 20 minutes to assess IP-1 accumulation over time. In some conditions, cells were pre-incubated with the P2Y1 antagonist MRS2500 for 10 minutes prior to ADP activation.

After incubation, 3 µL of IP-1-d2 conjugate and 3 µL of europium cryptate-labelled anti-IP-1 antibody, both dissolved in lysis buffer, were added to each well to lyse the cells. Following a 45-minute incubation at room temperature in the dark, time-resolved fluorescence was measured at 620 and 665 nm using a Tecan Spark multimode plate reader. HTRF ratios were converted to IP-1 concentrations in nM for unknown samples using a standard IP-1 curve provided by the manufacturer. Area under the curve (AUC) values were calculated and compared among conditions to assess differences in IP-1 levels in platelets.

**DNA constructs**

DNA constructs used for transient transfection of HEK293T cells for signalling profiling were either purchased online or created in-house. pcDNA3.1(+)/P2RY1 and pcDNA3.1(+)/P2RY12 were sourced form cDNA Resource Center (cDNA.org). pcDNA3.1/SRF-GrRluc, pcDNA3.1/2xSRE-Rluc, and pcDNA3.1/3xNFAT-Nluc were synthesised in house and will be described in more detail in Will et al., in preparation.

#### Cell culture and transfection

All experiments used human embryonic kidney SV-40 T antigen cells (HEK293T) gifted by Dr David Poyner (Aston University, UK). HEK293T cells were grown in DMEM/F-12 media (with or without pheonl red) with 1% AA solution (AA; Sigma-Aldrich) and 10% or 5% foetal bovine serum (FBS; Sigma-Aldrich) at 37 °C and 5% $CO_2$. For assays requiring overexpression of the receptor and other DNA components, transient transfection was conducted utilising a 6:1 v/w ratio of polyethyleneimine:DNA in sterile 150 mM NaCl. Transfection of cells occurred for 24 hours before replating and use in assays.

**G-CASE G protein dissociation assay**

Inhibition of G protein activation was completed using G-CASE plasmids (Cat #168121, 168125, 168127; Addgene).[5] HEK293T cells were transiently transfected via a 6:1 v/w ratio of polyethyleneimine:DNA in sterile 150 mM NaCl to express the receptor (400 ng) and G-CASE plasmid (400 ng) of interest. Cells were incubated in transfection media for 24 hours, then replated on opaque, white 96-well plates (Greiner) at 500 cells/µL and grown for 24 hours. Cell plates were incubated with 90µL of a HEPES-buffered solution (HBSS; 20 mM HEPES; 0.1% BSA; pH 7.3) with coelenterazine H (Nanolight Technology, USA) for 5 minutes without light. Cells plates were then stimulated with ligand and read using a PHERAstar® at 460nm and 528nm every 60 seconds for 15 minutes. Dose response curves were created by calculation of the BRET ratio and quantifying the area under the curve for the entire 15-minute read.

**Luminescence based genetic reporter assays**

For genetic reporter assays, HEK293T cells transfected via a 6:1 v/w ratio of polyethyleneimine:DNA in sterile 150 mM NaCl to express the receptor (1 µg) and genetic reporter plasmid (1 µg) of interest. Cells were incubated in transfection media for 24 hours, then replated on opaque, white 96-well plates (Greiner) at 1000 cells/µL and grown for 24 hours. During replating, cells transfected with either SRF or SRE constructs were resuspended in DMEM/F-12 media lacking phenol red dye and 0.5% FBS. Cells transfected with NFAT constructs were resuspended in DMEM/F-12 media lacking phenol red dye and

10% FBS. Ligands were diluted in assay media and applied and incubated with cells for 6 hours (SRF and SRE) or 4 hours (NFAT). Cells were then lysed with a lysing buffer containing 2mL deionized water, 0.5 mL 5x passive lysis buffer (Promega), and 5 µL of 5mM coelenterazine-H. Plates were shaken for 10 minutes away from light, then read with a BMG CLARIOstar® plate reader with no filter on the luminescence measurement and gain of 3600 RFU.

**Intracellular $Ca^{2+}$ mobilisation assays**

Intracellular $Ca^{2+}$ release ($[Ca^{2+}]_i$) was measured on untransfected HEK293T cells. Cells were plated on PLL coated, black, clear bottomed, 96 well plates (Greiner) at a concentration of 500 cells/µL and grown for 24 hours. Cell plates were incubated for 1 hour at room temperature with a solution containing 12 µL of 20% pluronic acid and 12 µL of Calbryte-520 (AAT Bioquest) in DMSO in 12 mL of HBS, followed by a wash and incubation in plain HBS for 45 minutes. Incubated cell plates were read with a FlexStation® 3 plate reader using SoftMax Pro 5.4 at 490 and 520 nm respectively. BAPTA was applied at a concentration of 5 nM for 30 seconds to chelate extracellular $Ca^{2+}$ before application of the ligands. Ligands were serially diluted in nominally free calcium buffer (NCF). Measurement of subsequent $Ca^{2+}$ release was taken for two minutes before a solution was applied to induce maximal release and florescence of unbound cytosolic dye (0.7% triton, 70mM $CaCl_2$ in HBS).

***In vivo* analysis of LPS-induced pulmonary inflammation**

All animal experiments were performed in accordance with the Animals (Scientific Procedures) Act 1986 with 2012 amendment, with local ethical approval from King's College London; and reporting herewith complying with the ARRIVE guidelines. Male and female BALB/c mice (7-12 weeks) were sourced from Charles River Laboratories Ltd. Litter mate male and female test/platelet $P2Y_{1-/-}$ mice (Homozygous for $P2Y_1$-*loxP* flanked allele and hemizygous for PF4-*cre)* and control 'wild type' mice (Homozygous for $P2Y_1$-*loxP* flanked allele and a non carrier for PF4-*cre*) were bred in house to test the efficacy of KSN159-27.[6] Animals were housed under standard SPF conditions of 22 ± 2 °C with a 12:12 light: dark cycle in cages of 4 mice. Animals were provided with food and water *ad libitum* and given environmental enrichment in the form of wood shavings, shredded paper, cardboard mouse houses and tubing.

**LPS induced lung inflammation in mice**

Groups of female Balb/c mice were treated with MRS2500 (3mg/kg), or KSN159-27 (0.01, 0.01, 1, 3, 10, 30mg/kg) or saline (vehicle) intravenously. 10 minutes post drug administration, mice were challenged with 1mg/kg LPS in 50µl via intranasal administration under isoflurane anaesthetic. 4 hours post LPS challenge, 1.5ml of bronchoalveolar lavage fluid was collected and processed for total and differential cell counts as previously described.[6,7] In studies using Plt-$P2Y_{1-/-}$ mice and litter mate controls, KSN159-27 was dosed at 10mg/kg, 10 minutes before, and 6 hours post LPS challenge before lavage at 24 hours. Circulating platelet numbers were enumerated at 4 hours post LPS challenge after blood was taken via cardiac puncture and quantified in stromatol (1:100 dilution) on an Improved Neubauer haemocytometer using an Axioskop Microscope under an x40 objective. To measure the bleeding time, a separate cohort of mice were kept under continuous anaesthesia with inhaled isoflurane anaesthetic. Bleeding assays were performed 1 hour post LPS challenge in mice by tail tip amputation, immersing the tail in saline at 37°C and continuously monitoring bleeding patterns. Each animal was monitored for up to 10 minutes and bleeding times determined using a stop clock. At the conclusion of the experiment, animals were killed with an overdose of anaesthetic.

**Measurement of thrombo-embolic responses**

Platelets from donor mice were prepared as previously described and resuspended in TAP buffer (10:1 mixture of CFTS and ACD with 2.5µM $PGE_1$).[8] Platelets were then incubated with 1.8 MBq $^{111}InCl_3$ in sterile 0.9% w/vol NaCL which was mixed 10 minutes earlier with 1mg/ml Tropolone at a 1:5 ratio. This produced lipophilic $^{111}In\text{-(tropolone)}_3$ complexes capable of penetrating cells membranes and dissociating into the cells. Radioactive counts were measured in samples at this point using a Single Point Extended Area Ratio (SPEAR) radiation detector with CAPture ™ technology. Platelets were then pelleted via centrifugation at 1500 g for 7 minutes and the radioactive supernatant removed. The platelet pellet was carefully washed then gently resuspended in CFTS to a volume of approximately 200µl per mouse. Radioactive counts were measured for a second time to calculate the percentage labelling efficiency in each preparation. Platelets were then left to rest for 45 minutes to allow the effects of $PGE_1$ on platelet function to dissipate. Radiolabelled platelets (1-1.5x$10^8$ platelets/mouse in 200µl) were infused intravenously via a lateral tail vein and returned to their cages. In order to assess platelet function *in vivo,* mice were anaesthetised using inhaled isoflurane in an induction box. Once anesthetised, mice were removed from the induction box and general anaesthesia maintained via nose cone inhalation. SPEAR radiation detectors with CAPture ™ technology were then placed over the pulmonary vasculature and platelet accumulation was non-invasively measured in real time following intravenous administration of either ADP or Up4A (5mg/kg) via a lateral tail vein following the production of a stable baseline reading. Samples were read until readings returned to baseline levels. The signal from the gamma detector was then deconvoluted using a UCS30 spectrophotometer and data logged on a computer running USX software. *In vivo* platelet aggregation/accumulation was then reported as a maximal percentage increase from the baseline reading or through calculation of the area under the curve (AUC) to calculate the overall size and duration of the response.

**Statistical analysis**

Data are expressed as mean ± SEM. Quantification of platelets via microscopy was conducted with the experimenter blinded to the sample identity. All other studies were quantified by machine (plate reader or flow cytometer). Chemotaxis and signalling activity data is normalised to a negative control to give a fold mean of control values, due to baseline variations between donors. Groups are of equal size and are indicated in figure legends. Power calculations were undertaken to provide an estimation of the minimum sample size to detect difference between two means, dependent on intra-group variability of assays based on previous published data.[3,4,7] In all studies, the group size indicated is the number of independent values, and the statistical analysis was therefore conducted using these independent values. All outliers are included in the data analysis and presentation. In studies where one-way ANOVA was used (as indicated in figure legends), with multigroup studies with parametric variables, post hoc tests (Dunnett's) were conducted only if F in ANOVA achieved *P* value of less than 0.05 and there was no significant variance in homogeneity. With subsequent post hoc tests, also a *P* value of less than 0.05 was considered significant.

### In vitro pharmacokinetic and toxicity assessment of KSN-159-27

Compound KSN-159-27 was evaluated in the *in vitro* pharmacokinetic study including solubility, absorption, and metabolism (**Table A**). For the solution properties, KSN-159-27 showed intermediate aqueous solubility of 143 µM in phosphate buffer solution (PBS) compared to reference compounds simvastatin and rifampicin. KSN-159-27 also showed high protein binding in human plasma, with 100% of the proteins bound, which is similar to that of the reference compounds sertraline (99%) and warfarin (99%).

The *in vitro* absorption of KSN-159-27 was assessed using the bi-directional permeability assay in Caco-2 cells. KSN-159-27 had a $P_{app}$ of $0.1x10^{-6}$ cm/s from apical side (A) to basolateral side (B) and a $P_{app}$ of $0.1x10^{-6}$ cm/s from B to A, resulted in an uptake ratio ($P_{app}$A-B/$P_{app}$B-A) of 1. The percent recovery was lower (18% A to B and 9% B to A) compared to the reference compounds in both directions (100% A-B and 93% B-A for propranolol, 100% A-B and 93% B-A for propranolol ), indicating potential problems like poor solubility or non-specific binding during the assay.

The *in vitro* metabolism of KSN-159-27 was assessed as its intrinsic clearance in human liver microsomes. KSN-159-27 had a shorter half-life (5.6min) than the reference compounds (terfenadine, 18.0 min and propranolol >120min). The resulted intrinsic clearance ($CL_{int}$) is 1234.2 µl/min/mg of microsome, which is much higher than terfenadine (385.8 µl/min/mg of microsome) and propanolol (<57.8 µl/min/mg of microsome). These indicated that KSN-159-27 has a relatively rapid metabolism in hepatic microsomes, therefore will cause less accumulation and potential toxicity.

**Table A. The *in vitro* ADME profile of compound KSN-159-27 and reference compounds.**

| Solution properties | | | |
|---|---|---|---|
| Aqueous solubility [a] | KSN-159-27 | Simvastatin | Rifampicin |
| In PBS, pH 7.4 (µM) | 145.1 | 30.1 | 200.0 |
| Protein binding [b] | KSN-159-27 | Sertraline | Warfarin |
| % Protein bound | 100 | 99 | 99 |
| % Recovery | 91 | 91 | 100 |
| ***In vitro* absorption** | | | |
| Permeability in Caco-2 cell [c] | KSN-159-27 | Propranolol | Labetalol |
| $P_{app}$A-B ($10^{-6}$ cm/s) | 0.1 | 29.6 | 5.2 |
| $P_{app}$B-A ($10^{-6}$ cm/s) | 0.1 | 39.6 | 31.3 |
| % Recovery (A-B) | 18 | 100 | 93 |
| % Recovery (B-A) | 9 | 93 | 93 |
| ***In vitro* metabolism** | | | |
| Intrinsic clearance (Human liver microsomes) [d] | KSN-159-27 | Terfenadine | Propranolol |
| $t_{1/2}$ (min) | 5.6 | 18.0 | >120 |
| $CL_{int}$ (µl/min/mg of microsomes) | 1234.2 | 385.5 | <57.8 |

[a] Aqueous solubility (µM) in PBS at pH 7.4 determined with high-performance liquid chromatography-ultraviolet spectroscopy (HPLC-UV). [b] Measure of percentage of protein binding and percentage of compound recovery during the assay determined in equilibrium dialysis using human plasma. [c] The permeability of compounds assessed in bidirectional Caco-2 cell permeability assay with pH = 6.5 for donor chamber and pH = 7.4 for receiver chamber. The extent of permeability is measured as apparent

permeability coefficient ($P_{app}$) from apical (A) to basolateral (B) or in reverse direction. The percentage recovery of compound is calculated as the total amount of compound in the donor and the receiver at the end of experiment/the amount of initial compound present. The uptake ratio of compound is calculated as $P_{app}$A-B/$P_{app}$B-A. [d] The metabolic stability of compounds was determined in 0.1mg/ml human liver microsomes, measured as the half-life ($t_{1/2}$) and apparent intrinsic clearance ($CL_{int}$).

The *in vitro* toxicity of KSN-159-27 was assessed using a hERG Potassium Channel Assay. (**Table B**) Results showing inhibition greater than 50% are considered to represent significant effects of test compounds. Compound KSN-159-17 gave an inhibition of 30.34% at 10 µM and therefore was not considered a significant liability, compared to reference compound verapamil that had an $IC_{50}$ of 0.38 µM.

**Table B. The *in vitro* toxicology profile of compound KSN-159-27 and reference compound verapamil.**

| *In vitro* CHO-hERG assay | | | |
|---|---|---|---|
| | Conc (µM) | % Inhibition | **IC50** (µM) |
| **KSN-159-27** | 10 | 30.34 | N/A |
| **Vehicle control** | | 6.85 | |
| **Verapamil** | 0.01 | 6.06 | 0.38 |
| | 0.03 | 19.15 | |
| | 0.1 | 22.47 | |
| | 0.3 | 38.83 | |
| | 1 | 74.29 | |
| | 3 | 89.38 | |

The parameters measured were the maximum tail current evoked on stepping to 40mV and ramping back to -80mV from the test pulse. All data were filtered for seal quality, seal drop, and current amplitude. The peak current amplitude was calculated before and after compound addition and the amount of block was assessed by dividing the Test compound current amplitude by the Control current amplitude. Control data is the mean hERG current amplitude collected 15 seconds at the end of the control period; Test compound data is the mean hERG current amplitude collected 15 seconds at the end of test concentration application for each concentration. KSN-159-27 tested in the presence of 0.1% Pluronic F-68 Non-Ionic Surfactant and at approximately room temperature. Vehicle is 0.33% DMSO Addition 1

#### Results

##### Supplementary Figure 1

Molecular Models showing 3D interaction of MRS2500, KMR-82-13 and KSN-159-27 with $P2Y_1R$

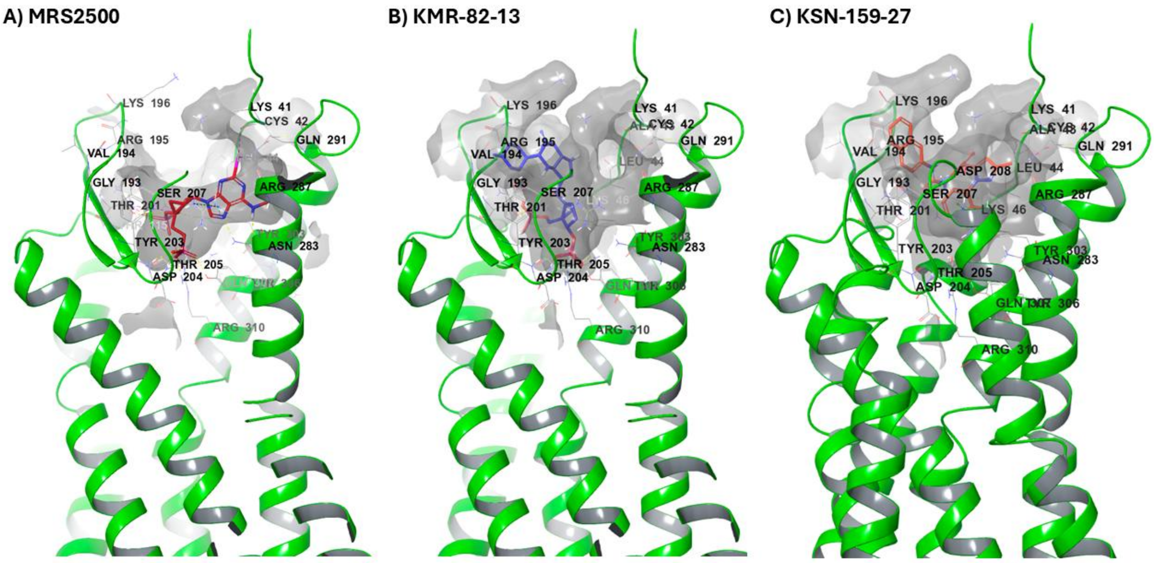

##### Supplementary Figure 2

A) Protein RMSF Analysis of P2Y1R During KSN-159-27 during the MD simulation

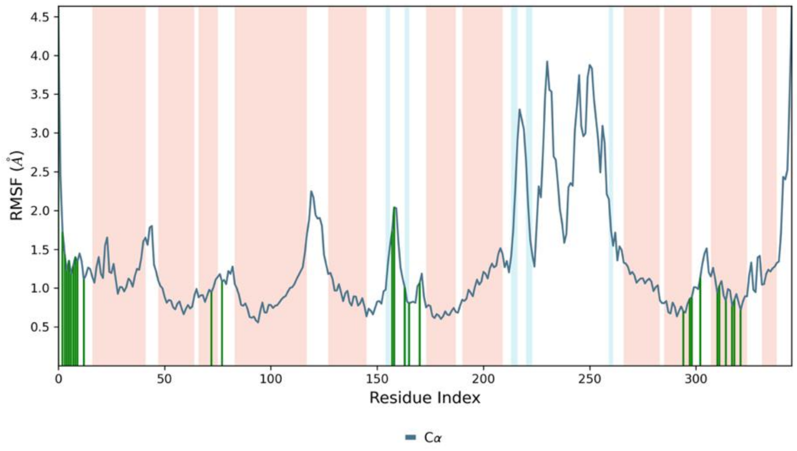

Protein RMSF analysis further supported stability of the ligand-binding region. The largest fluctuations were observed mainly in flexible loop or terminal regions outside the principal ligand-contacting pocket, including residues around 215-259 and the C-terminal region.

B) Secondary Structure Stability of P2Y1R During MD Simulation

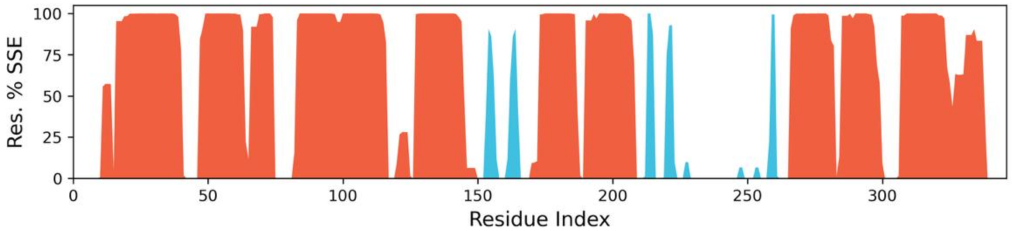

Secondary structure analysis showed that the overall helical and strand content of P2Y1R was maintained across the 200-ns trajectory, with total secondary structure content remaining approximately stable around 60%. Thus, KSN-159-27 binding did not induce detectable loss of receptor secondary-structure integrity during the simulation.

**Supplementary Figure 3**

2D Interaction Map of Synthesised Compounds with P2Y$_1$R

A) KSN 1-59-25

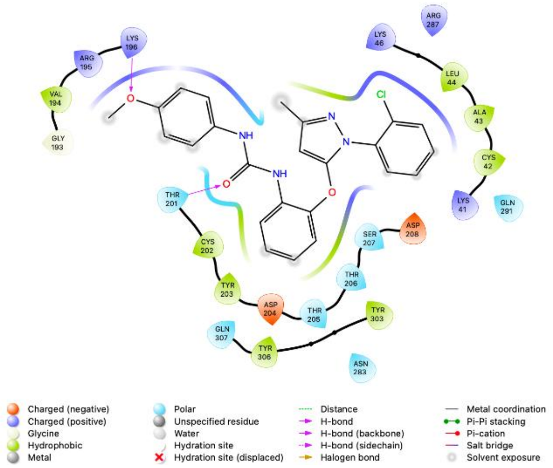

B) KSN159-26

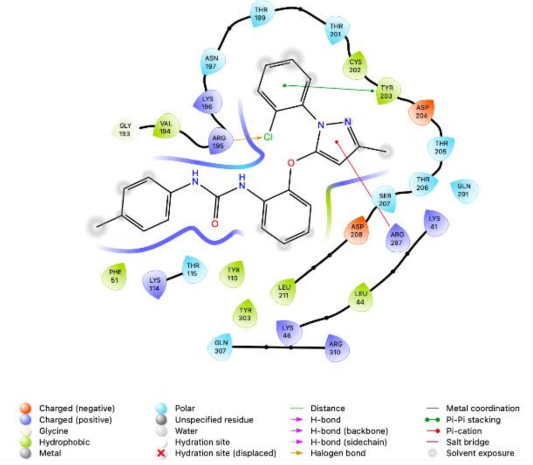

C) KSN-159-28

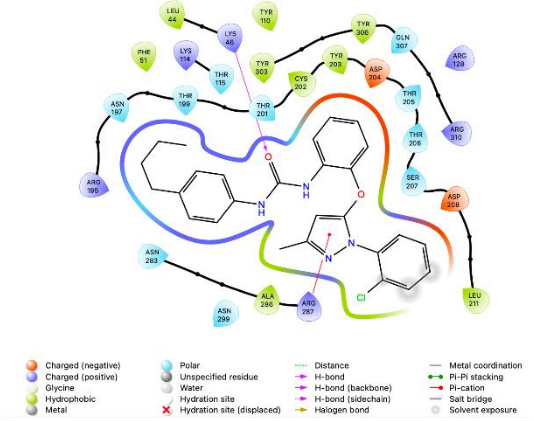

D) TA-167-129

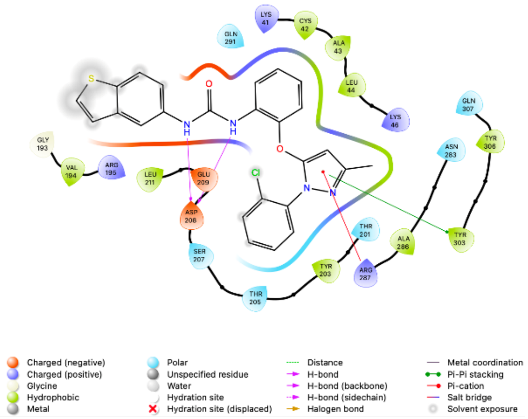

**Supplementary Figure 3 (contd.)** 2D Interaction Map of Synthesised Compounds with $P2Y_1R$

E) TA-167-137

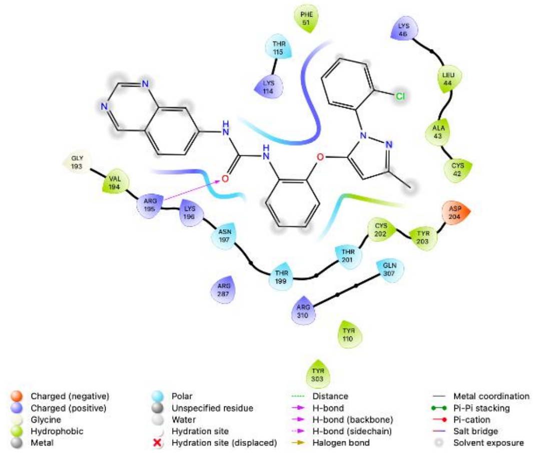

F) TA-167-139

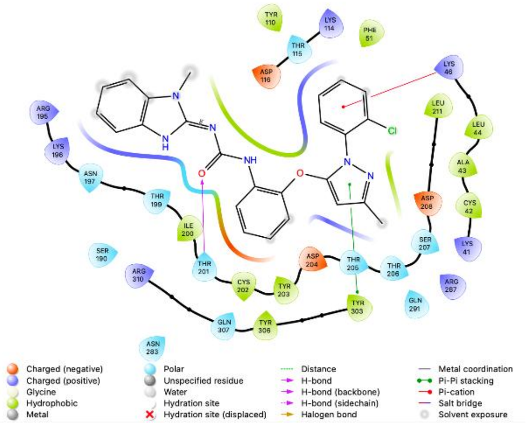

688

G) TA-167-153

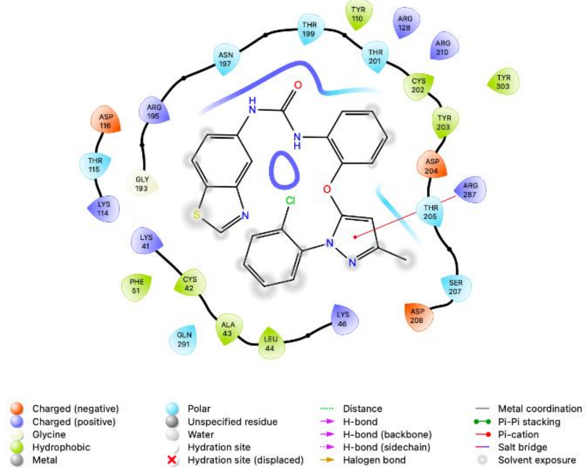

H) TA-167-155

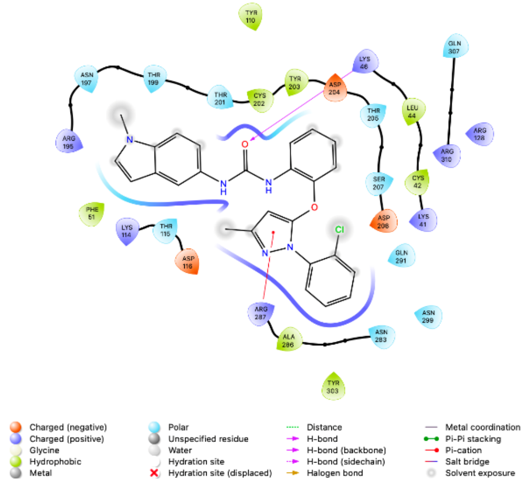

I) TA-167-157

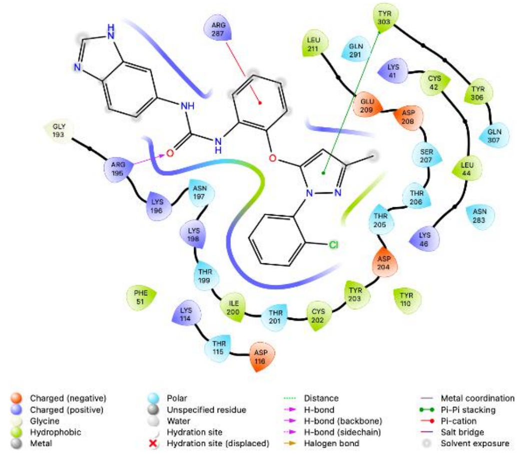

J) TA-167-161

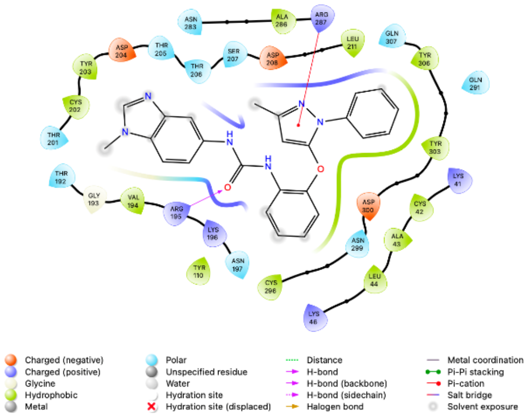

**Supplementary Figure 3 (contd.)**

2D Interaction Map of Synthesised Compounds with P2Y1R

K) TA-167-163

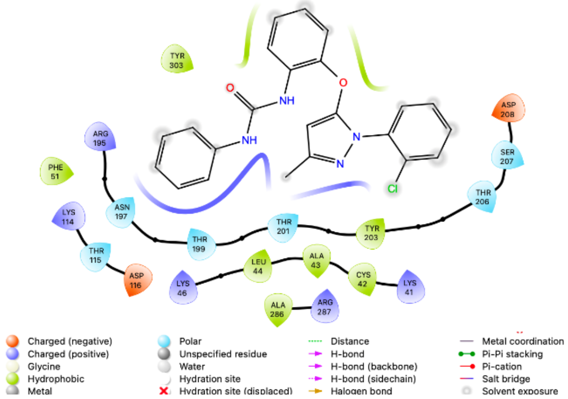

L) SL-183-5

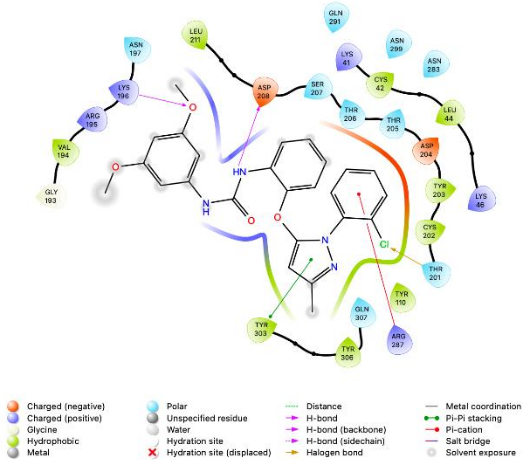

M) SL-183-6

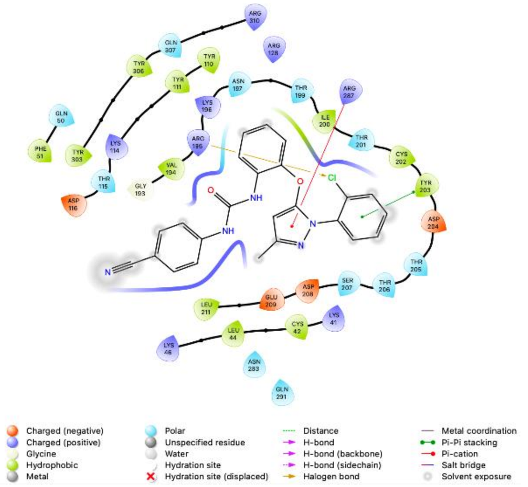

N) SL-183-7

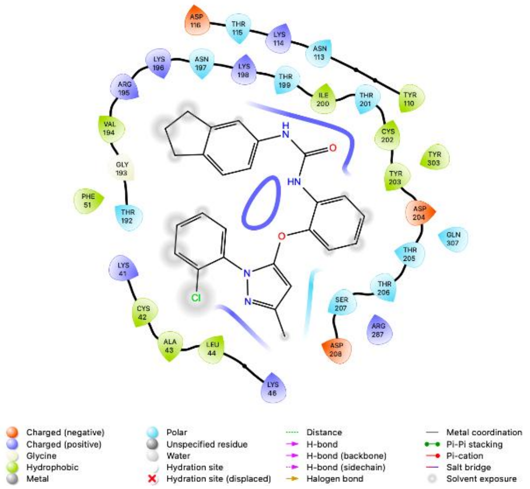

O) SL-183-8

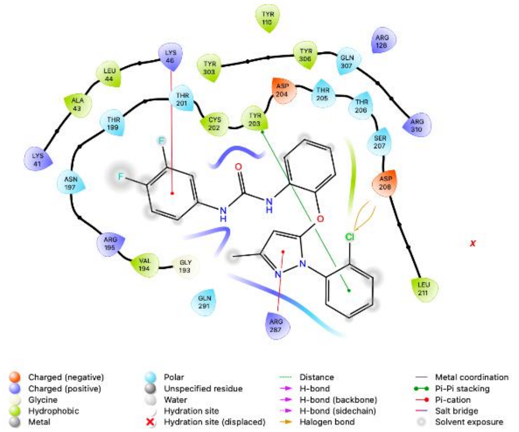

#### Supplementary Figure 4

##### A P2Y12R agonism: G protein dissociation

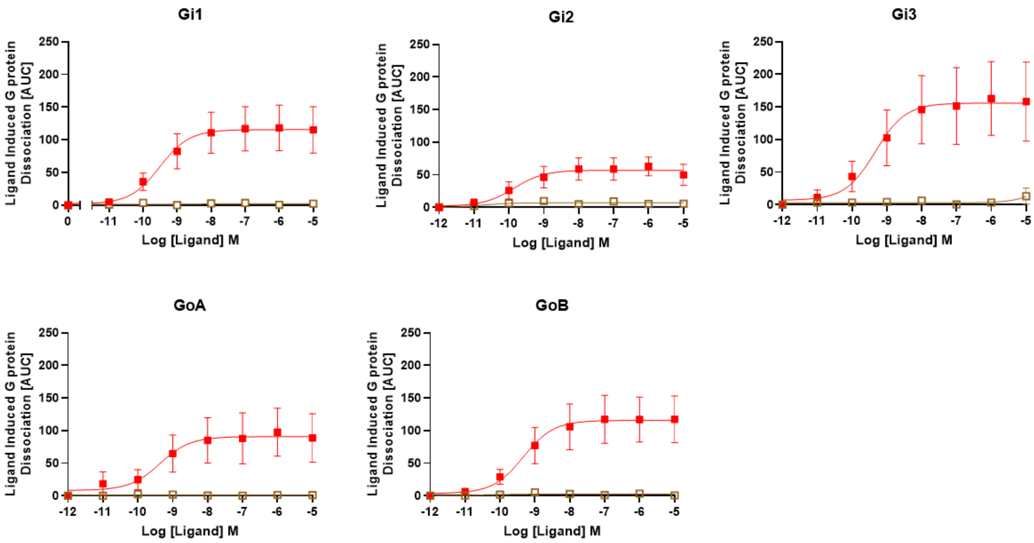

##### B P2Y12R agonism: Genetic reporter response

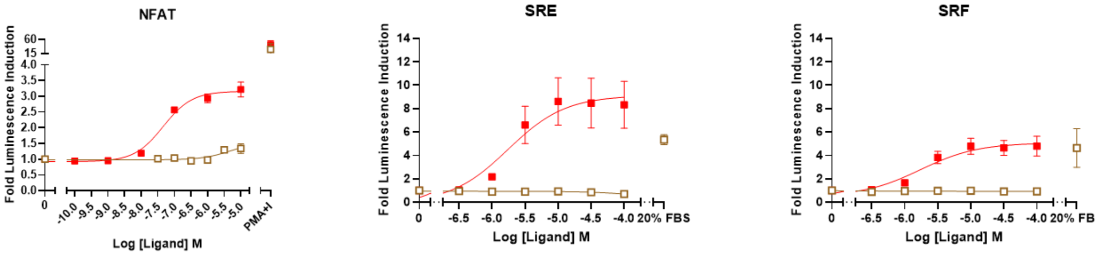

**Effect of KSN-159-27 on P2Y$_{12}$R activation.** HEK293T cells were transfected to express P2Y$_{12}$R and experiments were conducted to measure possible antagonistic effects of KSN159-29 (brown line) compared to 2MeSADP (red line) on: (A) G protein activity in using the Trupath assay towards G$_{i1}$ , G$_{i2}$ , G$_{i3}$, G$_{o}$A, and G$_{o}$B; and (B) using the genetic reporters NFAT-RE, SRE, and SRF.

#### Supplementary Figure 5

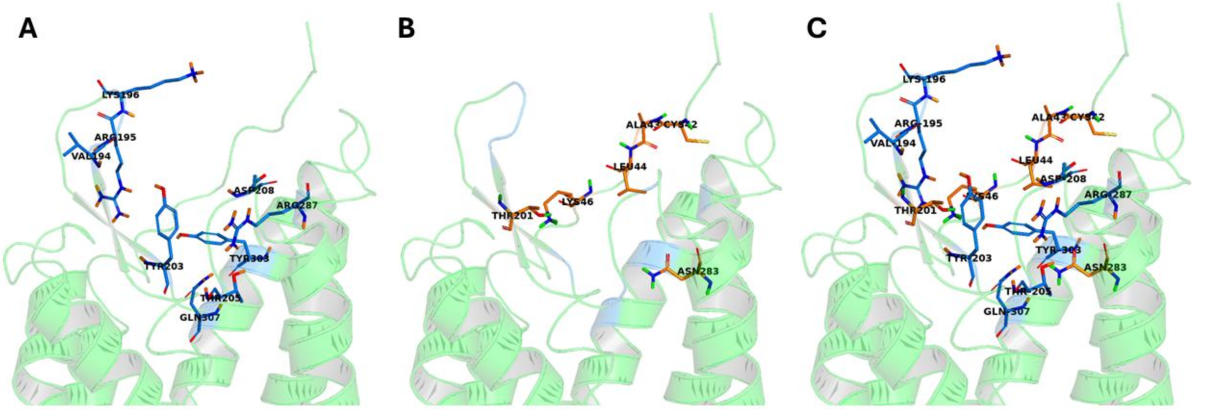

**3D Docking analysis of compounds that inhibit chemotaxis compared to compounds that do not inhibit chemotaxis.** Through *in silico* analysis, the residues that appear important for inhibiting chemotaxis are shown in marine blue (A), and the residues that appear not important in orange (B) and merged (C). Based on the available 2D interaction maps, the chemotaxis-inhibiting compounds (KMR-82-13, KSN-159-27, KSN-159-28, KSN-159-129, KSN-159-163) appear to share an extended binding orientation within the P2Y$_1$R orthosteric pocket, rather than a single conserved residue interaction. KSN-159-27 provides the clearest example of this pattern. Its hydrophobic naphthyl group appears to occupy a broad lipophilic region of the pocket, helping to stabilise an extended pose. KSN-159-28 may achieve a related effect through its butylphenyl group, although with different local anchoring. TA-167-129 also shows an organised interaction profile, including polar contacts around Asp208/Glu209 and π-type interactions around Arg287/Tyr303. TA-167-163 has fewer clear polar contacts but appears to retain similar hydrophobic pocket occupancy. In contrast, inactive compounds (TA-167-153, TA-167-161, SL-183-5) retain some individual contacts with residues such as Arg195, Lys196, Thr205, Asp208, Arg287 or Tyr303, but do not reproduce the same coordinated deeper-pocket binding pattern. For example, TA-167-153 and TA-167-161 engage Arg287 and/or Arg195, indicating that these interactions alone are insufficient. SL-183-5 forms several apparent contacts but remains inactive, further supporting the idea that interaction count alone does not predict activity. Overall, the emerging SAR hypothesis is that P2Y$_1$R-mediated chemotaxis inhibition requires a productive KMR-82-13/KSN-159-27-like geometry within the deeper orthosteric pocket. Compounds that preserve this binding geometry may be more likely to modulate chemotaxis-associated P2Y$_1$R signalling, whereas compounds with isolated or misoriented contacts may fail to produce a chemotaxis-selective inhibitory profile.

#### Supplementary Figure 6

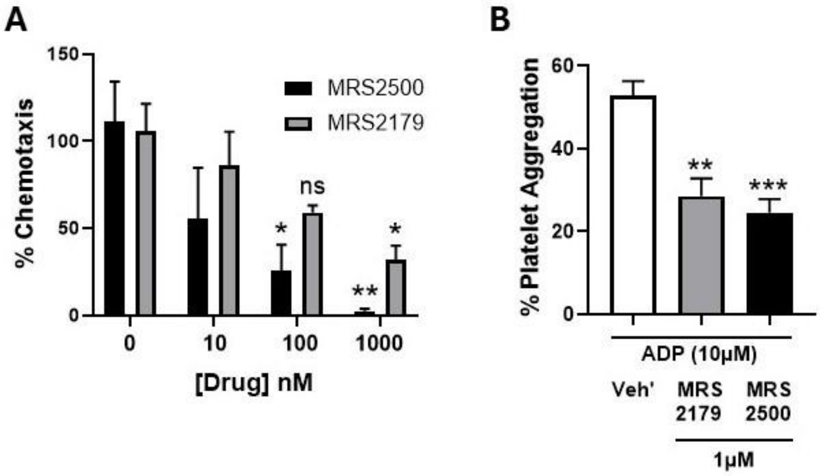

**Effect of MRS2179 on platelet functions in vitro.** Washed platelets from healthy human donors were incubated with $P2Y_1R$ antagonists MRS2179 or MRS2500 at concentrations stated and subjected to an inflammatory chemotaxis assay towards fMLP in the presence of ADP (100 nM) (**A**) or hemostatic aggregation assay induced by ADP (10 µM) (**B**). Data represented as means +/- SEM, n= 4 (**A**,**B**). Data analysed using two way ANOVA (**A**) or one way ANOVA (**B**) followed by Dunnett's test. *P<0.05, **P<0.01, ***P<0.001 vs vehicle control group. ns =not significant.

#### Supplementary Figure 7

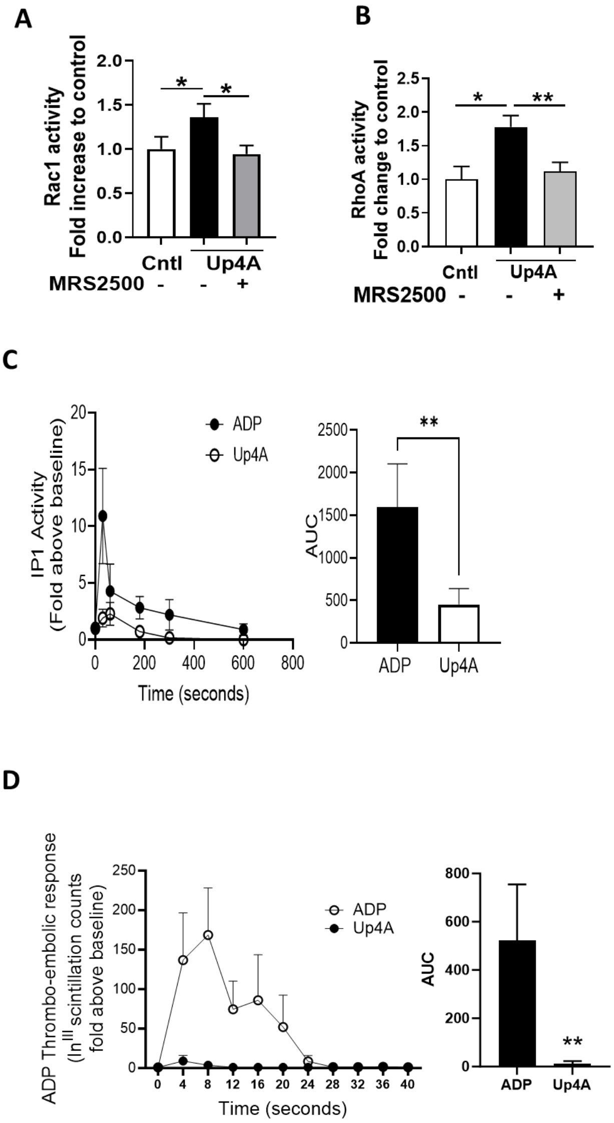

**Effect of Up4A on platelet P2Y$_1$ signalling events *in vitro* platelet thromboembolism *in vivo*.** Platelets from healthy human donors were incubated with 10µM Up4A to stimulate Rac1 (**A**) or RhoA activity (**B**); or IP1 production used as to measure PLC pathway (**C**). P2Y$_1$ antagonist MRS2500 (1uM) added where indicated 10 min before stimulation. Because previous *in vivo* data had shown Up4A was able to induce pulmonary platelet accumulation and inflammation when administered intranasally,[22] we used another *in vivo* model of platelet aggregation that causes acute (within a minute) platelet thromboembolism when ADP is administered i.v, to compare to Up4A administration (**D**). Thus, Up4A or ADP (5mg/kg) were administered to mice (i.v.) that had previously received In$^{111}$ labelled platelets, and scintillation counts were enumerated in the thorax as a measure of thromboembolism (**D**). Data is means +/- SEM. *<0.05, **<0.01. n =6 (**A-C**), 7 (**D**).

#### Supplementary Figure 8

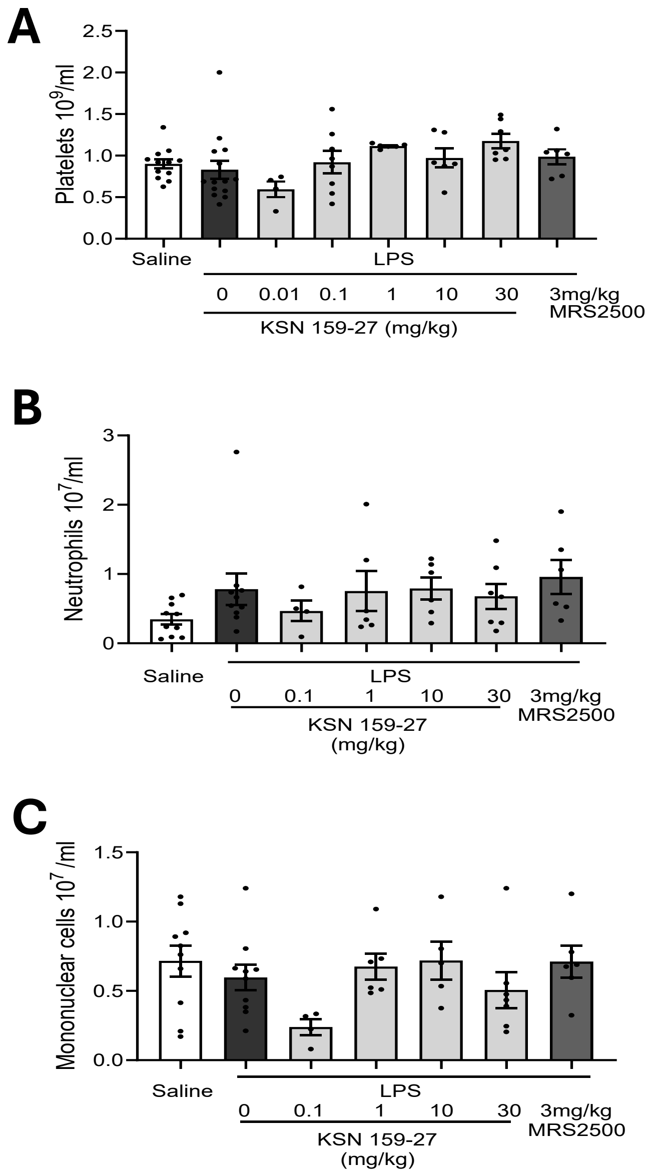

**Circulating platelet and leukocyte numbers in mice administered KSN-159-27.** Studies were conducted in BALB/C mice to establish whether the intravenous administration of KSN-159-27 (0.01, 0.1, 1, 10, 30 mg/kg) or MRS2500 (3mg/kg) 5-10 minutes before LPS exposure (1mg/kg intranasal). Tail bleeds were performed 4 hours after LPS exposure in anaesthetised mice and platelets (**A**) neutrophils (**B**) and mononuclear cells (**C**) were enumerated. Means +/- SEM n= 4-10 as indicated for groups (A-C) because data was amalgamated from 4 separate studies to study dose-response.
